# The RNA-binding protein RBP45D of Arabidopis plays a role in epigenetic control of flowering time and DCL3-independent RNA-directed DNA methylation

**DOI:** 10.1101/2021.06.22.449407

**Authors:** Liangsheng Wang, Duorong Xu, Kristin Habermann, Wolfgang Frank, Dario Leister, Tatjana Kleine

## Abstract

RNA-directed DNA methylation (RdDM) helps to defend plants against invasive nucleic acids. In the canonical form of RdDM, 24-nt small interfering RNAs (siRNAs) are produced by DICER-LIKE 3 (DCL3). Here, we describe the *Arabidopsis thaliana prors1* (*LUC*) transgenic system, in which transcriptional gene silencing (TGS) is independent of DLC3. A forward genetics screen performed with this system identified both known components of RdDM, and the RNA-binding protein RBP45D. RBP45D promotes DNA methylation, and its loss delays flowering, especially at high temperature, presumably mediated by elevated FLC levels. RBP45D is localized to the nucleus, where it is associated with snRNAs and snoRNAs. RBP45D maintains siRNA production originating from the *LUC* transgene, but does not alter mRNA levels or affect processing of transcripts of known RdDM genes. We suggest that RBPD45 facilitates DCL3-independent siRNA production by stabilising either the precursor RNA or the – as yet unidentified – slicer protein.

## Introduction

In plants, invasive nucleic acids of viral, transposon or transgenic origin, but also endogenous genes, can be transcriptionally silenced by RNA-directed DNA methylation (RdDM) (Gebert and Rosenkranz, 2015; Matzke and Mosher, 2014; Pooggin, 2013; Wendte and Pikaard, 2017; Zhou and Law, 2015). RdDM is also involved in several aspects of development. For example, it modulates flowering time, participates in biotic and abiotic stress responses including the heat shock response, and can silence transposable elements (TEs) that could otherwise undergo transposition under heat stress (Erdmann and Picard, 2020; Zhang et al., 2018). According to the current view, the canonical RdDM pathway is initiated by RNA polymerase IV (POL IV)- mediated transcription, which produces short single-stranded RNAs (ssRNAs) of roughly 30 to 45 nucleotides in length (Blevins et al., 2015). RNA-DEPENDENT RNA POLYMERASE 2 (RDR2) converts these into double-stranded RNAs (dsRNAs), which are then cleaved by DICER-LIKE 3 (DCL3) into 23- and 24-nucleotide (nt) small interfering RNAs (siRNAs) (Singh et al., 2019). The 24-nt siRNAs are loaded onto ARGONAUTE (AGO) proteins, mainly AGO4 and AGO6, and pair with complementary scaffold RNAs, which are produced by POL V. The AGO-siRNA-Pol V complex then recruits chromatin-modifying enzymes, including the DNA methyltransferase DOMAINS REARRANGED METHYLASE 2 (DRM2), which catalyzes *de novo* DNA methylation in a sequence-independent manner (Zhang et al., 2018). Following DNA replication, CG, CHG and CHH (where H is A, C or T) methylation are maintained by METHYLTRANSFERASE 1 (MET1), CHROMOMETHYLASE 3 (CMT3), and CMT2, respectively (Wendte and Pikaard, 2017). Maintenance of CHH methylation through RdDM depends on DRM1 and DRM2 (Matzke et al., 2015; Stroud et al., 2013; Wendte and Pikaard, 2017; Zhang et al., 2018). DNA methylation can be reversed by active demethylation, which is initiated by DNA demethylases, of which REPRESSOR OF SILENCING 1 (ROS1) is the major representative in *Arabidopsis thaliana* (Zhu, 2009).

Transgenes are widely used to study gene function and regulation, and to introduce novel and desirable properties in plant-breeding campaigns. Transgene silencing (TGS) by RdDM and other mechanisms has therefore proved problematic for plant researchers. On the other hand, efforts to understand how transgenes are silenced have ultimately revealed much of what we know about the RdDM pathway (Erdmann and Picard, 2020). One prominent system employs the *ros1* (*RD29A-LUC*) transgene. Mutations in *ROS1* cause transcriptional silencing of both the transgenic, *RD29A* promoter-driven, luciferase reporter gene (*RD29A-LUC*) and the endogenous *RD29A* gene (Gong et al., 2002). Screening for second-site suppressors of *ros1* mutant plants has in turn identified many RdDM components, including RDM2 (RNA-directed DNA methylation 2, NRPD4, NRPE4), which is a component of Pol IV and Pol V complexes (He et al., 2009).

Numerous questions about the RdDM pathway remain unanswered, among them whether DCL-independent routes of siRNA biogenesis exist or not (Singh and Pikaard, 2019; Yang et al., 2016; Ye et al., 2016; Zhang et al., 2018).

Here, we introduce the *Arabidopsis thaliana prors1* (*LUC*) transgene system, which enables 24-nt siRNAs to be generated independently of DCL3. We conducted a second-site mutation screen with the *prors1* (*LUC*) mutant to identify suppressor mutants in which TGS is reversed. Among already known components of RdDM, we identified the RNA-binding protein RBP45D (AT5G19350). Col-0 plants lacking RBP45D were then generated using CRISPR/Cas, and whole-genome bisulfite sequencing (WGBS) of these lines showed that RBP45D is involved in changes in DNA methylation. Loss of RBP45D also results in a late flowering phenotype especially under heat stress, presumably mediated by the release of epigenetic repression of FLC levels. RNA-Seq analysis showed that RBP45D does not boost mRNA levels, and has no influence on the processing of transcripts of known RdDM genes. The protein is localized in the nucleus, where it is associated with snRNAs and snoRNAs. Small RNA-Seq and Northern analyses revealed that RBP45D promotes the production of siRNAs originating from the transgene. Our results indicate that RBP45D is an active component of a DCL3-independent RdDM pathway.

## Results

### The *prors1* (*LUC*) mutant provides a new probe for silencing components

Insufficiency of the prolyl-tRNA synthetase PRORS1, which is targeted to both plastids and mitochondria, results in impaired organellar gene expression (OGE), activates retrograde signalling and alters expression levels of nuclear genes encoding chloroplast proteins like the light-harvesting chlorophyll a/b binding protein Lhcb1.2 (Leister and Kleine, 2016; Pesaresi et al., 2006). In order to systematically screen for factors that are involved in OGE-dependent signalling, we first introduced into Col-0 plants a reporter construct that allows the luciferase (*LUC*) gene to be expressed under control of the *LHCB1*.*2* promoter (Col-0 (*LUC*); Figure 1A). Col-0 (*LUC*) was then crossed with *prors1-2* to generate *prors1* (*LUC*) plants. The latter displayed much lower LUC activity than Col-0 (*LUC*) plants (Figure 1B). Measurements of *LHCB1*.*2* and *LUC* RNAs confirmed the reduction of *LHCB1*.*2* mRNA levels in *prors1* (*LUC*) to 36% of that in Col-0 and Col-0 (*LUC*). Strikingly, *LUC* levels fell to 3% of the Col-0 (*LUC*) value (Figure 1C), which might be caused by hypermethylation in the context of TGS (Matzke et al., 2000). To test whether the reduced *LUC* levels in *prors1* (*LUC*) were indeed due to high methylation levels, Col-0, *Col-0* (*LUC*) and *prors1* (*LUC*) were grown on MS medium alone or on MS medium containing 5-aza-2’
s-deoxycytidine (5Aza-dC), a chemical inhibitor of DNA methyltransferase (DNMT) activity that should reduce methylation levels. Indeed, *prors1* (*LUC*) seedlings treated with 5Aza-dC displayed much higher LUC activity than those grown under control conditions (Figure 1D), comparable to that in the Col-0 (*LUC*) strain grown on control MS plates. The same was true of 3-week-old plants that had been sprayed daily with 5Aza-dC during the third week (Figure 1E). Because no *PRORS1*-unrelated mutation could be identified that might have activated silencing of foreign transgenes in the *prors1* (*LUC*) mutant (Table S1), all T-DNA insertion sites were mapped by resequencing of *prors1* (*LUC*) plants (Table S1). In addition to the insertion of the *LUC* construct into the 3′ region of *AT5G47160*, the construct was found in the intron region of *AT5G12400* which, like the *PRORS1* locus *AT5G52520*, lies on chromosome 5. Given the close proximity of these insertions, the second insertion might have gone unnoticed in previous TAIL-PCR and segregation experiments, but it could explain *LUC* repression by silencing of transgenes through *trans*-inactivation (Daxinger et al., 2008; Matzke et al., 2000).

**Figure 1.**
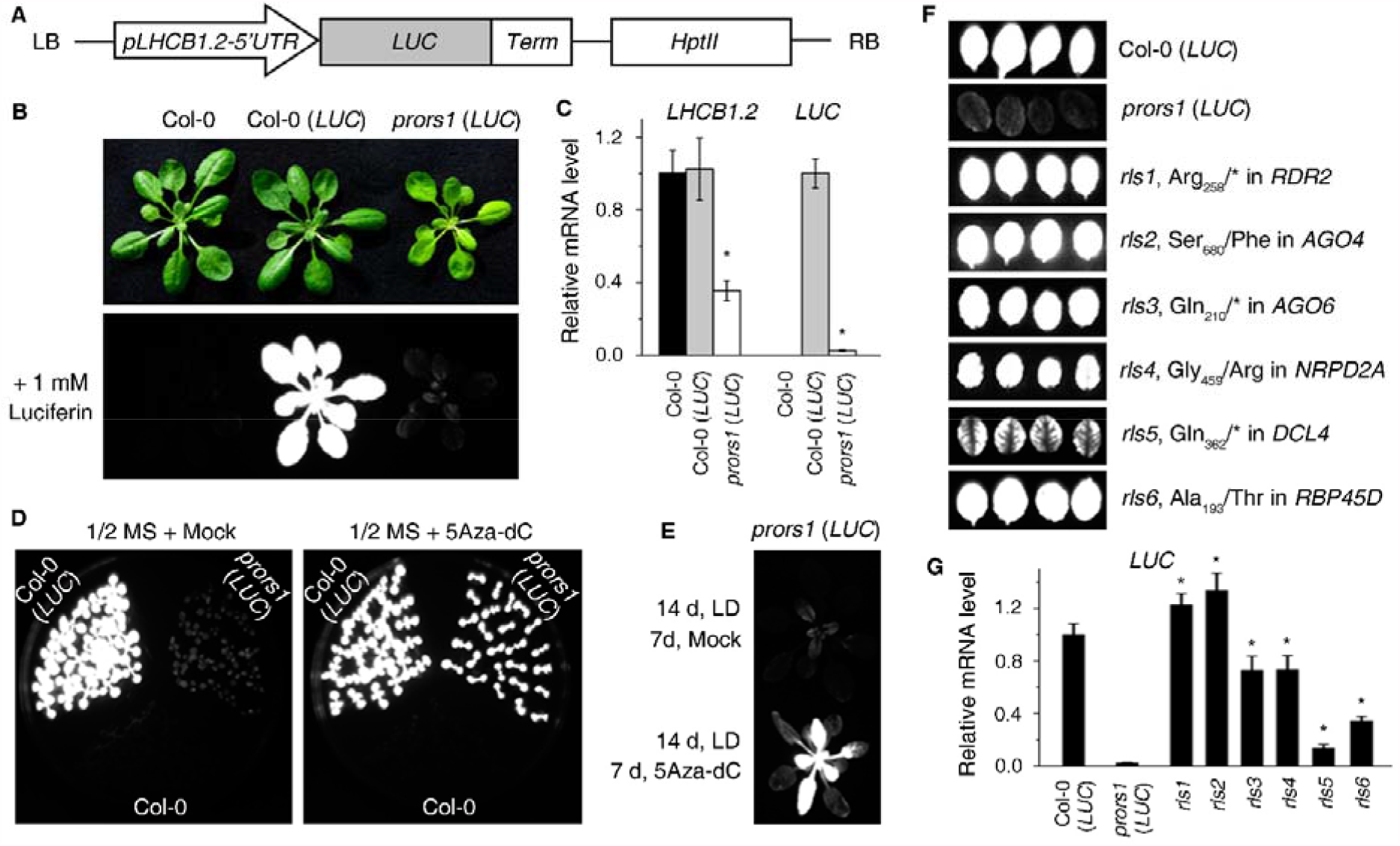
Identification of Arabidopsis *rls* (*<u>r</u>elease of <u>L</u>HCB1*.*2 <u>s</u>uppression*) mutants. See also Figure S1, Tables S1 and S2. **(A)** Schematic representation of the reporter construct used in this study. *HPTII*, hygromycin phosphotransferase II; LB, left border; *LUC*, coding region of the luciferase gene; *pLHCB1*.*2- 5’
sUTR*, promoter and 5’ UTR region of the *LHCB1*.*2* gene; Term, terminator. **(B)** Expression of the *LHCB1*.*2*:*LUC* transgene in Col-0, Col-0 (*LUC*) and the *prors1* (*LUC*) mutant. Plants were grown in long-day (LD; 16 h light/8 h dark) conditions for 3 weeks (upper panel) and were then sprayed with 1 mM luciferin to detect LUC activity (lower panel). Col-0 was used as a negative control. **(C)** Relative *LHCB1*.*2* and *LUC* mRNA levels in 3-week-old Col-0, Col-0 (*LUC*) and *prors1* (*LUC*) plants were determined by quantitative RT-PCR analysis. AT4G36800, encoding an RUB1-conjugating enzyme (RCE1) was used as a control. The RNA expression levels are reported relative to those in Col-0 (*LHCB1*.*2* mRNA levels) or Col-0 (*LUC* mRNA levels), which were set to 1. Data are represented as mean values from three independent experiments each with three replicates. Bars indicate standard deviations (SDs). Significant differences were evaluated with Student’s t-test (p < 0.05) and are denoted by asterisks (*). **(D)** LUC activity in 7-day-old seedlings grown in the absence (mock) or presence of the DNA methylation inhibitor 5Aza-dC. **(E)** LUC activity in plants grown for 14 days without inhibitor and of 3-week-old plants that were sprayed daily with 5Aza-dC in the third week. **(F)** LUC phenotypes and corresponding mutation sites in identified *rls* mutants. **(G)** Relative *LUC* mRNA levels in *rls* mutants. Quantitative RT-PCR analysis was conducted as in **(C)**. Data are mean values of three independent measurements with their SDs. Asterisks indicate significant differences (*P* < 0.05, *t*-test) between *rls* mutants and *prors1* (*LUC*).

In spite of this, the screen was pursued; *prors1* (*LUC*) seeds were mutagenized with ethyl methanesulfonate (EMS) and the M2 generation was screened for altered LUC expression in 26-day-old plants. This approach resulted in the identification of putative suppressor mutations of the *prors1* (*LUC*) phenotype, which were named “<u>r</u>elease of *<u>L</u>HCB* <u>s</u>uppression” or *rls* mutants. The mutations in the recessive *rls* mutants were identified by a next-generation sequencing approach employing single-nucleotide polymorphism (SNP) ratio plotting (Etherington et al., 2014). Four of the six identified genes – *NRPD2A, RDR2, AGO4* and *AGO6* – code for proteins involved in the canonical RdDM pathway, and *DCL4* encodes a component of the post-transcriptional gene silencing (PTGS) and non-canonical RdDM pathways (Erdmann and Picard, 2020; Figure 1F, Table S2). Quantitative RT-PCR confirmed that *LUC* mRNA levels in the *rls* mutants were higher than in *prors1* (*LUC*). The smallest boost in *LUC* mRNA levels was observed for *rls5* (a *prors1* (*LUC*) *dcl4* double mutant; Figure 1G) and is reflected in the LUC activity in *rls5* plants (Figure 1F). The identified mutations were verified by crossing *prors1* (*LUC*) plants with corresponding T-DNA lines and observing a high LUC phenotype and/or by complementation of the respective *rls* mutant and obtaining a low LUC phenotype (Figure S1, Table S2). Note that the introduction of mutant alleles of *NRPD1* (*nrpd1- 3*; RNA polymerase IV subunit), *NRPE* (*nrpe1-11*; subunit of RNA polymerase V) and *DDM1* (*ddm1-10*; a SWI2/SNF2-like chromatin remodeling factor) into *prors1* (*LUC*) also restored *LUC* expression. Crosses between *prors1* (*LUC*) and *rdr1-6, rdr6-15, dcl2-1, dcl3-1, ago1-37, ago7-1* or *ago9-2* had little or no effect (Figure S1A).

In addition to known components of silencing pathways, the mutation of alanine 193 to threonine in RNA-binding protein 45D (RBP45D, UniProtKB: BQ8VXZ9) restored *LUC* expression in *rls6* (Figure 1F). The RBP45D gene *AT5G19350* is also close to *PRORS1* (which precludes verification of the mutation by crossing). However, LUC activity in *rls6* was suppressed by the introduction of RBP45D-Myc (and to a lesser extent RBP45D-YFP) into *rls6* (Figure S1B), suggesting that RBP45D is a silencing factor. To test whether the *rls6* suppression phenotype was caused by demethylation of *LHCB1*.*2*:*LUC*, 26-day-old *Col-0* (*LUC*), *prors1* (*LUC*), *rls6* and – as representatives of the RdDM pathway – *prors1* (*LUC*) *ago4-6* and *prors1* (*LUC*) *rdr2-2*, were subjected to whole-genome bisulfite sequencing (WGBS). While the *LHCB1*.*2*:*LUC* region in Col-0 (*LUC*) was not methylated, both the *LHCB1*.*2* promoter and the 5′ UTR region were hypermethylated in all sequence contexts (CG, CHG and CHH) in *prors1* (*LUC*) (Figure 2A), confirming that *LUC* silencing indeed results from hypermethylation of DNA. Hypermethylation was substantially decreased by introducing *AGO4* or *RDR2* mutations into *prors1* (*LUC*) (Figure 2A), which is consistent with the notion that the RdDM pathway induces methylation of a DNA template that leads to TGS (Erdmann and Picard, 2020). At first sight, the overall methylation rate of the transgene in *rls6* appeared to be the same (CG) or even higher than in *prors1* (*LUC*) (Figure 2A and B). But closer inspection of the methylation plot revealed reduced methylation of the proximal promoter and 5′ UTR region of *LHCB1*.*2* (Figure 2A), which was confirmed by chop-PCR (Figure 2C). Moreover, a plot of the methylation differences between *rls6* and *prors1* (*LUC*) across the whole *LUC* cassette reveals regions of higher and lower CG, CHG or CHH methylation of *rls6* compared to *prors1* (*LUC*) (Figure 2D).

**Figure 2.**
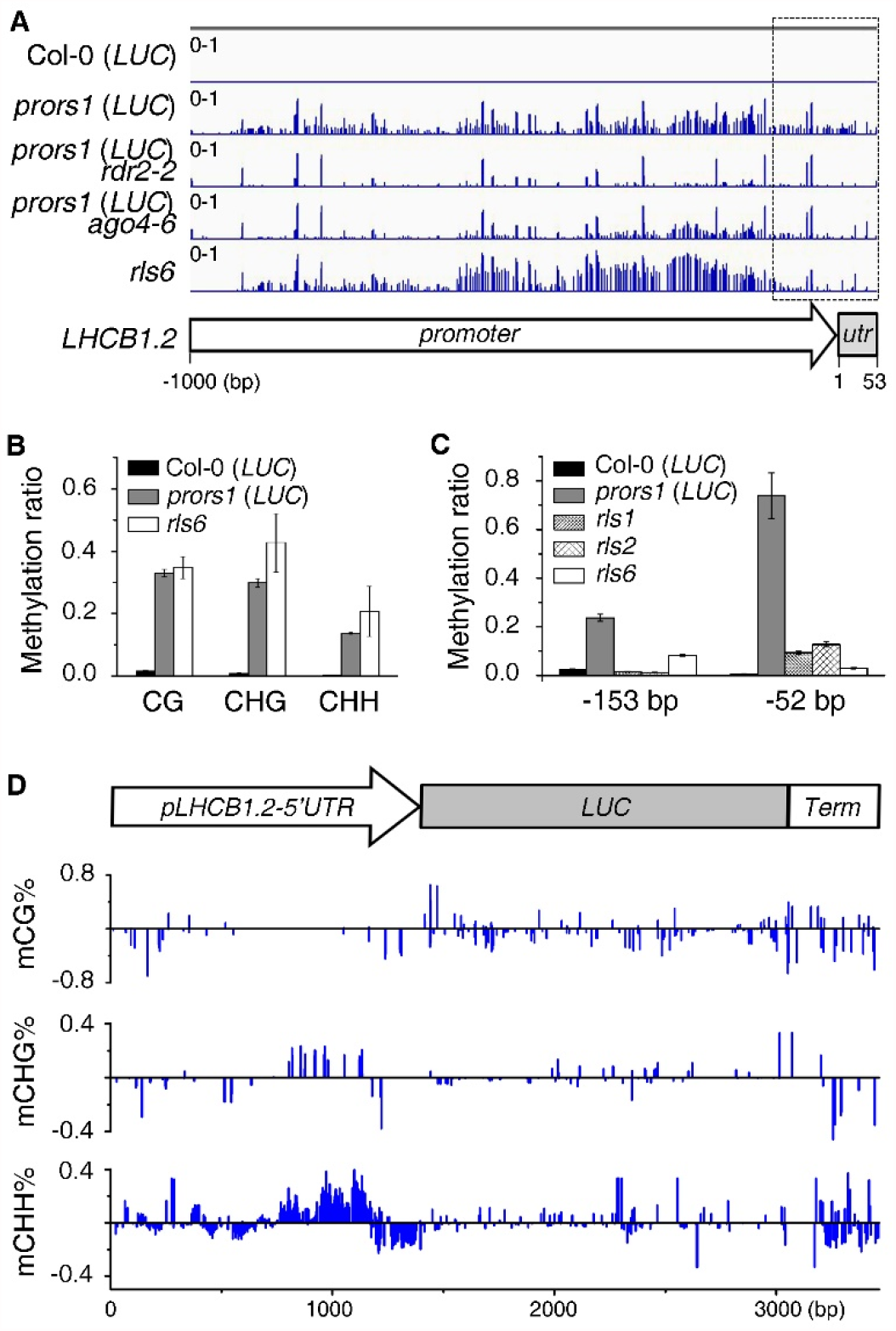
Characterization of the *LHCB1*.*2*:*LUC* transgene methylation status in the *rls6* mutant. **(A)** Whole-genome bisulfite sequencing was performed to detect changes in DNA methylation in the indicated mutants. Methylation levels of the *LHCB1*.*2* promoter and 5’ UTR were visualized with the Integrative Genomics Viewer (IGV). Note that the methylation level of the 3′ part of the *LHCB1*.*2* promoter and the 5′ UTR (marked by a dashed rectangle) are reduced (relative to *prors1* (*LUC*)) in the *prors1* (*LUC*) *rdr2-2* (reconstruction of the *rls1* mutant), *prors1* (*LUC*) *ago4-6* (reconstruction of the *rls2* mutant) and *rls6* mutants. **(B)** Graph displaying the overall CG, CHG and CHH levels in the transgene. Values represent mean values of three independent replicates with their SDs. **(C)** Confirmation of the reduced methylation level of the region marked by the dashed rectangle in panel **(A)** using chop-PCR. The numbers represent the positions at the *LHCB1*.*2* promoter, see also panel **(A)**. **(D)** Relative CG, CHG and CHH methylation levels of the *LHCB1*.*2*:*LUC* transgene in *rls6* compared to *prors1 (LUC)*.

Thus, although the screen failed to identify components of OGE-dependent retrograde signalling, it is capable of pinpointing elements of silencing pathways, such as RBP45D.

### Functional impairment of RBP45D affects DNA methylation

To investigate the function of RBP45D in a wild-type (WT) background, two T-DNA insertion lines (GK-746D12 and SAIL_1304H10) were identified in the SIGnAL database (http://signal.salk.edu/cgi-bin/tdnaexpress). Because we could not detect any T-DNA insertion in SAIL_1304, and GK-746D12 did not abrogate the expression of the full-length *RBP45D* mRNA, four independent CRISPR/Cas-mediated *rpb45d* knock-out lines (*rbp-ko1* to -*4*) were generated (Figure 3A). Subsequently, whole-genome bisulfite sequencing (WGBS) of Col-0 and the *rbp-ko1* line was conducted to gain a broader understanding of how loss of RBP45D affects the DNA methylome. We identified 5,447 differentially methylated regions (DMRs; differential methylation >0.1), of which 4,477 and 970 regions were hypo- or hypermethylated, respectively (Figure 3B, Table S3). Overall, promoters of protein-coding genes (PCGs) tended to be CG and CHH hypomethylated, and transposable elements (TEs) CHG and CHH hypomethylated in *rbp-ko1* (Figure 3B). The distribution of DNA methylation along gene bodies, TEs, and their 1-kb upstream and downstream flanking sequences revealed decreases in CG and CHH methylation, but no change in CHG methylation, in gene bodies compared to Col-0 (Figure 3C). CHG and CHH methylation of TEs were slightly and significantly reduced, respectively, while CG methylation was unchanged (Figure 3C). The hypomethylation of 5S rDNA in all sequence contexts, and of 66 genes whose gene products are involved in the defense response, is particularly noteworthy (Table S3), as mutants that are impaired in histone de- acetylation, the RdDM pathway or in DNA methylation maintenance typically exhibit diminished 5S rDNA methylation (Vaillant et al., 2007), and the defense response is epigenetically controlled (Espinas et al., 2016). These findings, together with the fact that its loss affects overall methylation status, point to a role for RBP45D in epigenetic silencing.

**Figure 3.**
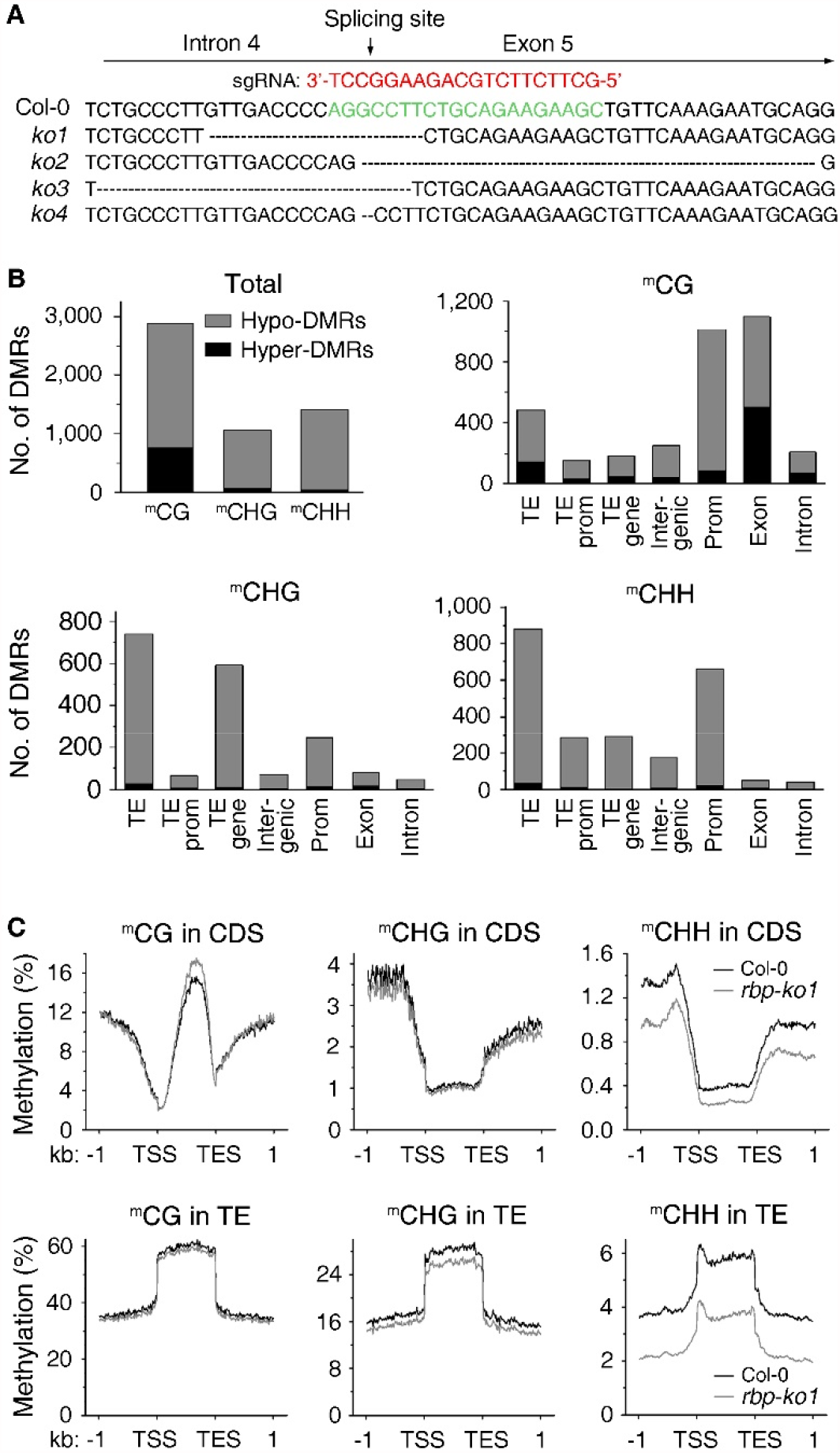
Loss of RBP45D affects DNA methylation. See also Table S3. **(A)** Generation of RBP45D knockout lines (*rbp-ko*) using the CRISPR/Cas9 technology. The single guide RNA (sgRNA) corresponding to the 4^th^ intron and 5^th^ exon of the RBP45D gene is shown in red letters. The DNA sequence changes in four independent *rpb45d* knockout lines (*ko1* to *ko4*) as determined by Sanger sequencing are also shown. **(B)** Whole-genome methylation analysis of the *rbp-ko1* mutant. Graphs display the number of differential methylated regions (DMRs) compared to Col-0. Intergenic, intergenic region; prom, promoter; TE, transposable element. **(C)** Plots of average CG, CHG and CHH methylation in coding sequences (CDS) and transposable elements (TE), respectively, and their 1-kb flanking regions. TES, transcription end site; TSS, transcription start site.

### RBP45D promotes flowering at high temperature

Five of the eight genes encoding proteins involved in the control of flowering time were either hypo- or hypermethylated, respectively, in *rbp-ko1* relative to Col-0 (Table S3). Several of these proteins act on FLOWERING LOCUS C (FLC), a repressor of flowering, which is known to be regulated by methylation (Sheldon et al., 1999): (i) METHYL-CPG-BINDING DOMAIN 9 (MBD9) is involved in modulating the acetylation state of *FLC* chromatin, which affects *FLC* expression levels (Peng et al., 2006), (ii) mRNA CAP-BINDING PROTEIN 80 (CBP80; ABA HYPERSENSITIVE 1, ABH) participates in processing of mRNAs for several floral regulators, including FLC (Kuhn et al., 2007), (iii) UDP-GLUCOSYL TRANSFERASE 87A2 (UGT87A2) controls flowering time via FLC (Wang et al., 2012), and (iv) VERNALIZATION INDEPENDENCE 4 (VIP4) is a positive regulator of *FLC* expression (Zhang and van Nocker, 2002). As *FLC* mRNA levels were elevated in *rbp-ko* lines (Figure S2A), their growth phenotypes and flowering times were investigated. Under long-day (LD) growth conditions, the *rbp-ko* lines showed no obvious growth or developmental defects (Figure 4A), and flowering was only slightly delayed (Figure 4B). However, it was significantly delayed under short-day (SD) growth conditions (Figure S2B). Moreover, RBP45D is annotated as being involved in the “response to heat” in the TAIR database (https://www.arabidopsis.org/), the RdDM pathway participates in basic heat-stress tolerance by mediating DNA methylation changes (Popova et al., 2013), and FLC is a repressor of temperature-induced flowering (Gan et al., 2014). Therefore, *rbp-ko* lines were grown for two weeks under LD conditions at 23°C, then transferred for 18 days to 32°C LD conditions. This regime substantially delayed flowering and fresh weight was reduced (Figure 4C, D, E).

**Figure 4.**
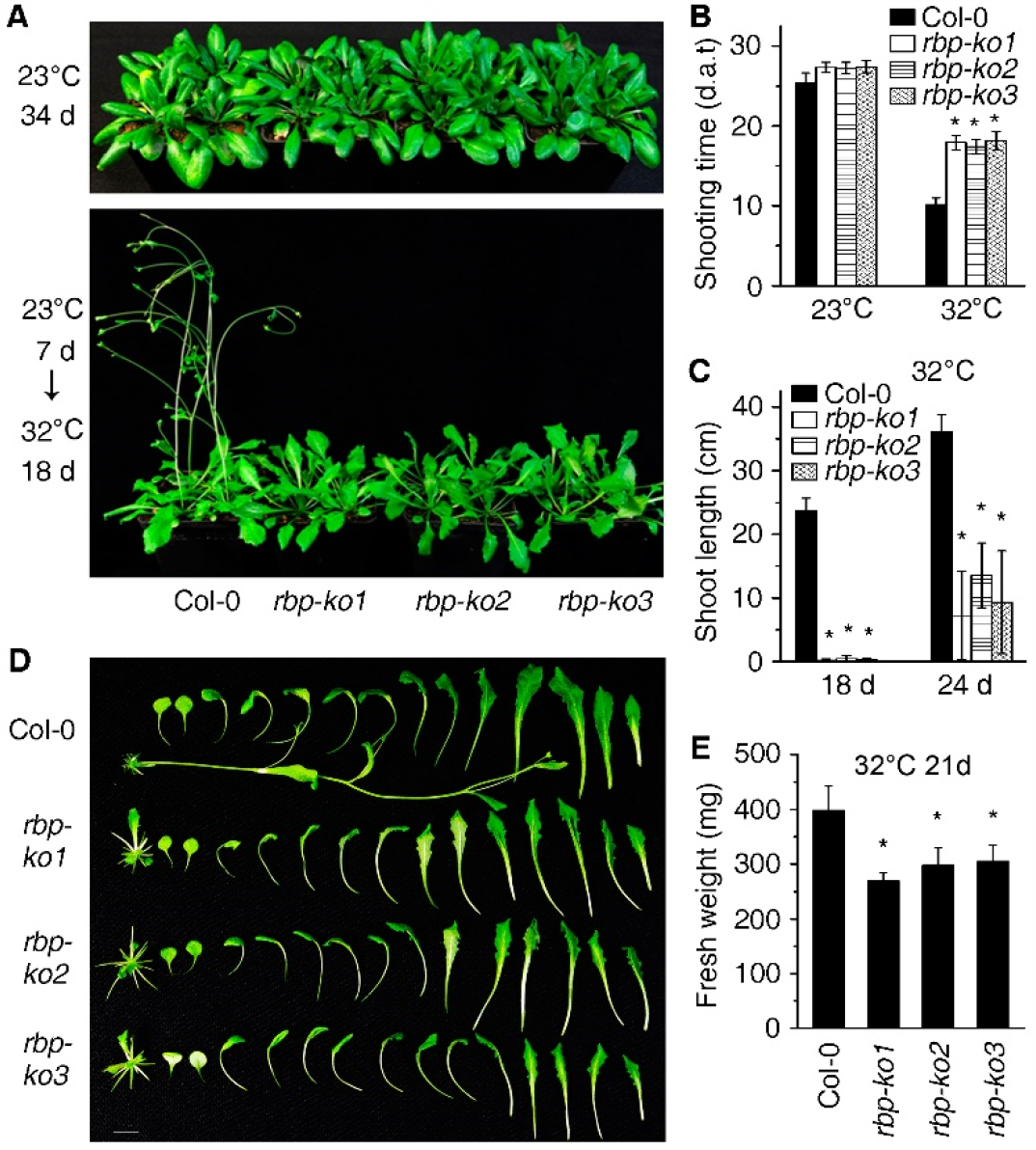
Loss of RBP45D promotes flowering at high temperature. See also Figure S2. **(A)** Flowering phenotypes of the three *rpb45d* knockout lines *rbp-ko1* to -*3*. Plants were grown in long-day (LD; 16 h light/8 h dark) conditions either for 34 days at 23°C (upper panel) or for 7 days at 23°C and were then shifted for 18 days to 32°C (lower panel). **(B, C)** Quantification of shooting times **(B)** and shoot lengths **(C)** of the three *rbp-ko* lines grown under optimal (23°C) or heat-stress (32°C) conditions. Data are shown as mean values (±SD) from three different plant pools. Each pool contained more than 12 plants. Significant differences (*P* < 0.05; *t*-test) with respect to Col-0 are indicated by asterisks. Note that the days are counted by day after transfer (d.a.t) following 7 days under normal growth conditions. **(D, E)** Close-up views **(D)** and fresh weights **(E)** of Col-0 and *rbp-ko* plants that were first grown for 7 days at 23°C and then transferred for 21 days to 32°C. Data represent mean values (±SD) of three independent experiments, each conducted with at least 12 plants. Asterisks indicate significant differences (*P* < 0.05, *t*-test) with respect to the Col-0 control.

Delayed flowering of *rbp-ko* lines under elevated temperature is presumably caused by elevated FLC levels owing to methylation changes, which again implicates RBP45D in RdDM.

### RBP45D does not promote or splice mRNA expression of genes related to the RdDM

To explore whether transcriptional levels of endogenous genes and TEs were altered by loss of RBP45D, 26-day-old Col-0 and *rbp*-*ko1* and -*2* lines were subjected to mRNA sequencing (mRNA-Seq) and those that changed by more than 2-fold were identified. Including *RBP45D* itself (Figure S3A), 181 and 496 transcripts were reduced and elevated, respectively, in both knock-out lines (Figure 5A, Table S4), and these two gene sets (“up” and “down”) were subjected to further analysis. Gene Ontology (GO) analysis identified in the set of “up” genes the biological process (BP) terms “systemic acquired resistance”, “response to virus” and “response to bacterium” as 11-, 9- and 7-fold significantly enriched, respectively (Figure 5B). In both gene sets “defense response” was flagged as 4-fold enriched, but there was no correlation between these genes and the hypomethylated defense-related genes (Figure S3B). Transcripts of 7 and 41 TEs were reduced and elevated, respectively, in both knock-out lines compared to Col-0 (Figure 5C, Table S4), suggesting that more TEs were released than silenced. However, the apparent release of TE expression was predominantly due to TE regions that overlapped with upregulated transcripts, such as the *PATHOGENESIS-RELATED GENE 1* (*PR1*) which was 12-fold up-regulated (Figure 5D). Moreover, seven genes associated with the function of FLC, including *FLC* itself (confirming qRT-PCR data, see Figure S2A), were up-regulated, but here again no correlation could be established with the FLC-associated genes that were differentially methylated (Figure S3B).

**Figure 5.**
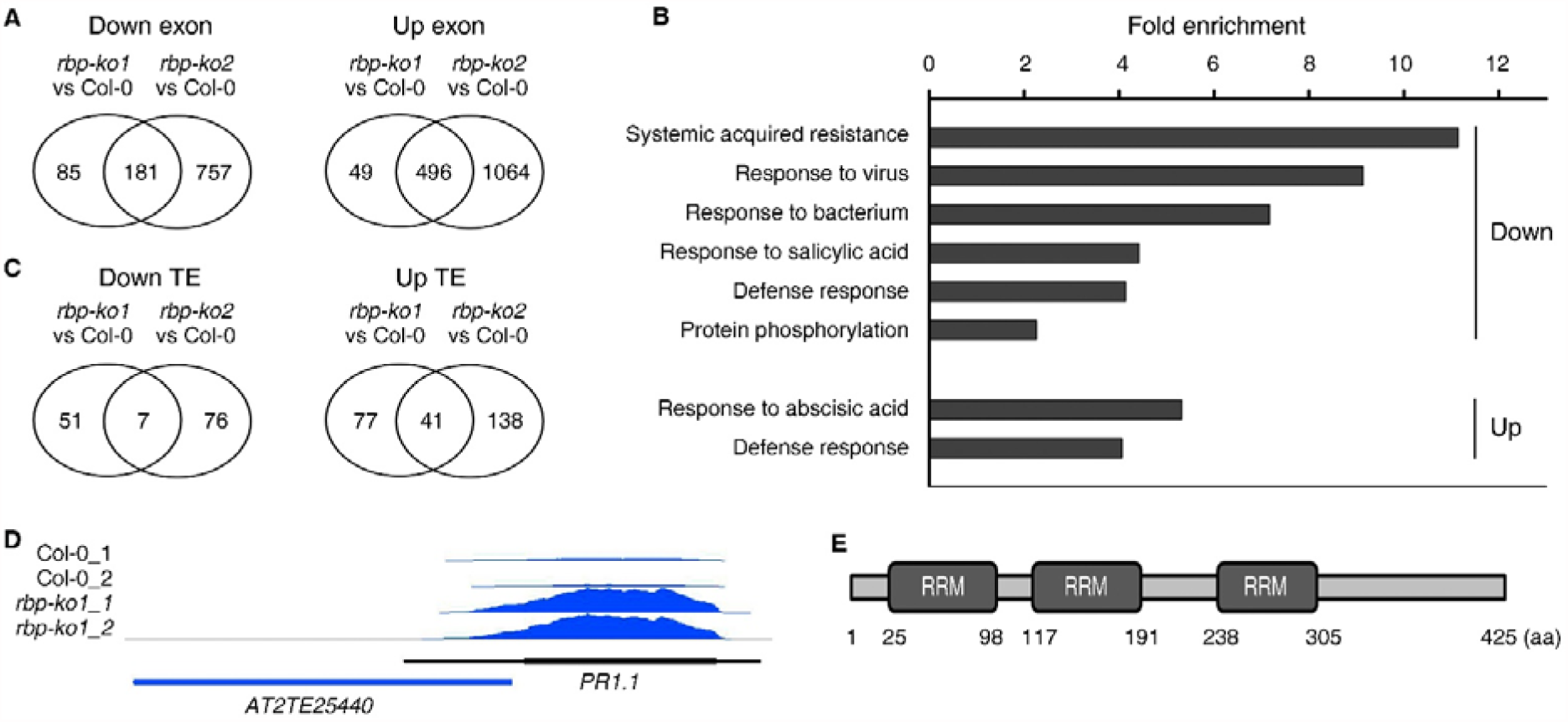
RBP45D regulates the expression of defense-related genes. See also Figures S3, S4 and S5 and Tables S4 and S5. **(A)** Numbers of genes whose expression is reduced (down) or elevated (up) more than 2-fold in *rbp-ko1* and *rbp-ko2* mutant lines compared with Col-0 grown for 26 days in LD conditions. **(B)** Gene Ontology (GO) analysis of genes affected in both *rbp-ko* mutants, i.e. the overlap between the gene sets shown in **(A)**. GO annotations for the biological process category were extracted from DAVID (Huang et al., 2007). GO terms with a >2-fold change and Benjamini corrected value of <0.05 are shown. **(C)** Numbers of TEs whose expression is reduced (down) or elevated (up) more than 2-fold in *rbp-ko1* and *rbp-ko2* mutant lines compared with Col-0. **(D)** IGV snapshot of the *PR1* gene region that overlaps with a transposable element. **(E)** Scheme illustrating the three predicted RNA recognition motifs (RRM) of RBP45D. The amino acid (aa) numbers are indicated.

RBP45D contains three RNA-recognition-motif (RRM) domains (Figure 5E). Accordingly, RBP45D has been identified as a poly(A) RNA- or mRNA-binding protein in RNA-binding proteome studies (Bach-Pages et al., 2020; Reichel et al., 2016). Hence, the function of RBP45D in silencing could be indirect; e.g., it might bind to or alter expression of mRNAs for proteins involved in silencing. We therefore assessed the degree of overlap between the “up” and “down” gene lists and a list of genes whose products are involved in epigenetic processes (Wang et al., 2020b). No overlap was found with the “down” gene set, while eleven genes from the “up” gene set were identified (Figure S3C), including genes for DCL3, DCL4, RDR6 and NRPB1 (the largest subunit of RNA polymerase II), but increased activity of these in the *rls6* mutant would not explain its silencing phenotype.

During the course of this work, RBP45D was identified in a forward genetics screen using a green fluorescence protein (GFP) splicing-reporter system (Kanno et al., 2020). The U1 small nuclear ribonucleoprotein (snRNP) factor PRP39A, which is involved in alternative splicing (Kanno et al., 2017), and the aforementioned CBP80 (Kanno et al., 2020) were also identified in that screen. Interestingly, levels of small RNAs derived from the *CBP80* locus are strongly reduced (see Table S7) and CBP80 is hypermethylated (Table S3) when RBP45D is lacking. Moreover, PRP39A is listed as a putative interactor with RBP45D in plant.MAP, which predicts interactions based on co-fractionation mass spectrometry (McWhite et al., 2020). Analysis of splicing in *rbp-ko1* revealed slightly elevated splicing at AGGCT and ATCT junctions, and reduced splicing at AGGG junctions (Table S5). We also detected exon-skipping and intron-retention events (Figures S4 and S5). Interestingly, these occurred in genes for splicing factors and proteins involved in flowering, like the flowering repressor FLOWERING LOCUS M (FLM; Figure S4). Remarkably, PRP39A was differentially spliced in the absence of RBP45D (Figure S4). Apart from the DCL1-4 proteins, RNASE THREE LIKE (RTL) proteins contain the RNASE THREE slicer domain. We detected slightly enhanced numbers of reads mapping to intron regions of the RNASE THREE LIKE *RTL1* (in *rbp-ko2* but not *rbp- ko1*) and *RTL2* (Figure S5A), but this was not consistent across replicates, and was not supported by qRT-PCR analysis (Figure S5B). No other splicing defect was detected in genes encoding RdDM components.

Together, these results indicate that transcripts for components of the RdDM are slightly elevated, but correctly spliced in *rbp-ko* lines. Therefore, RBP45D does not indirectly influence TGS through the expression or splicing of genes coding for RdDM components.

### RBP45D is a nuclear protein that binds to snRNAs and snoRNAs

Assuming a direct involvement of RBP45D in RdDM, we asked where it acts in the pathway, which encompasses steps in the nucleus and the cytosol (see Graphical Abstract; Gebert and Rosenkranz, 2015). RBP45D is predicted to be localized in the cytosol (https://suba.live; Hooper et al., 2017) and/or the nucleus (https://www.arabidopsis.org/). Confocal microscopy detected RBP45D in the nuclear matrix in roots of stably transformed *35S*:*RBP45D-YFP* (Col- 0) plants (Figure 6A). Western blotting of nuclear and cytosolic fractions prepared from protoplasts confirmed a predominantly nuclear localization, and showed that a small amount of RBP45D is present in the cytosol (Figure 6B).

**Figure 6.**
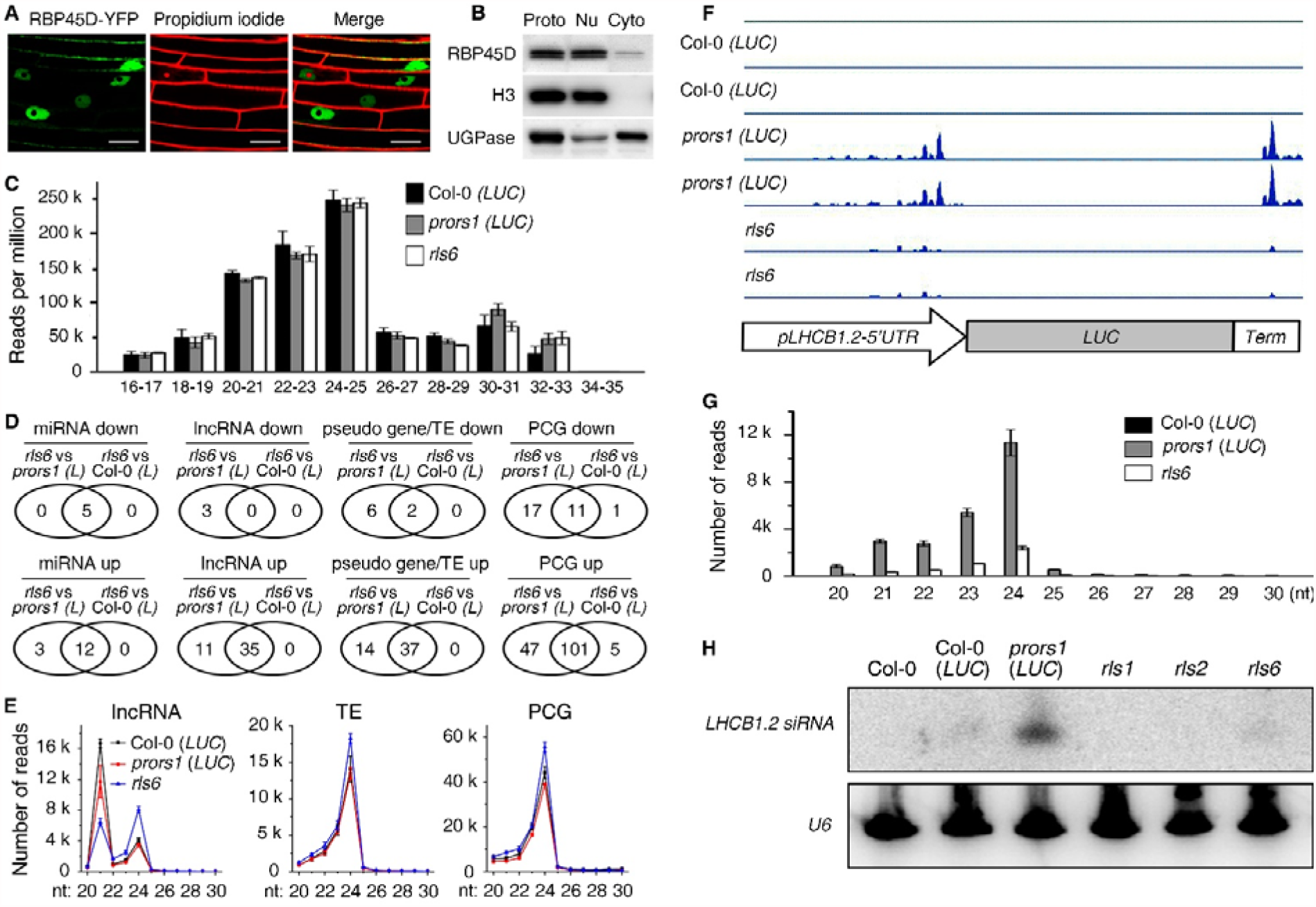
RBP45D promotes the generation of siRNAs originating from the *LHCB1*.*2*:*LUC* transgene. See also Figure S6. **(A)** RBP45D is localized predominantly in the nuclear matrix. Fluorescence microscopy of root tips from 6-day-old *35S*:*RBP45D-YFP* transgenic seedlings. Root tips were excised and stained with propidium iodide before confocal images were taken. Bar = 20 μm. **(B)** Confirmation of RBP45D localization by Western-blot analysis of purified protoplasts (Proto) which were fractionated into nuclei (Nu) and cytosol (Cyto). Exposure to antibodies against H3 and UGPase served as controls for the enrichment of nuclear and cytosolic proteins, respectively. **(C)** Comparison of the overall abundance and size distribution of small RNAs (sRNA) in Col- 0, *prors1 (LUC)* and *rls6* as detected by sRNA-Seq. **(D)** Venn diagrams showing the numbers of sRNAs corresponding to miRNAs, lncRNAs, pseudo genes/TEs, and PCGs that were increased (up) or decreased (down) more than 2-fold in *rls6* compared to *prors1 (LUC*) or Col-0 (*LUC*), respectively. **(E)** Size distribution plots of the relative abundance of siRNAs originating from lncRNA, TEs and PCGs in Col-0 (*LUC*), *prors1* (*LUC*) and *rls6*. **(F)** Read depths of siRNAs mapping to the *LHCB1*.*2*:*LUC* transgene, visualized with the IGV tool. **(G)** Graph displaying the numbers and lengths of reads mapping to the *LHCB1*.*2* promoter and 5’
s UTR region. **(H)** Northern-blot analysis illustrating the abundance of one prominent *LHCB1*.*2* siRNA species. Probing of U6 served as a loading control.

To identify the targets of RBP45D, RNA immunoprecipitation experiments were performed on *rls6* mutants complemented by expression of Myc-tagged RPB45D (see Figure S1B). Libraries prepared from RNAs that co-immunoprecipitated with RBP45D-Myc [and another unrelated protein-Myc fusion (Lee et al., 2021) as a control] were subjected to RNA sequencing (RIP-Seq). Based on the functions of its orthologs in other organisms, RBP45D has been suggested to be a component of the U1 snRNP (Kanno et al., 2020). Our enrichment analysis relative to the control experimentally confirmed the association of RBP45D with snRNPs. The small nuclear RNA (snRNA) U5.10 was the second most enriched target (66-fold enrichment, Table 1, Table S6), and other snRNAs identified belonged to the U2 snRNP family (U2.2 to U2.7, 9- to 30-fold enrichment), while snRNA U1A was only 3-fold enriched (Table S6). The small nucleolar RNAs (snoRNAs) AtsnoR6-3, -R6-2, -U14b, -R12-1a, and -R106, *FRUCTOSE-BISPHOSPHATE ALDOLASE 5* (*FBA5*) and one novel transcribed region were also identified as prominently enriched targets (>50-fold enrichment, Table 1, Table S6). With an enrichment of 20- to 47-fold, RNAs corresponding to 37 loci coding for proteins (none of them involved in RdDM), four snoRNAs and one snRNA, were among the intermediately enriched RNAs relative to the prominently enriched targets (Table S6). Notably, there was no significant overlap of RIP targets with differentially expressed loci identified in sRNA- and mRNA-Seq experiments (Figure S6A, B).

**Table 1.**
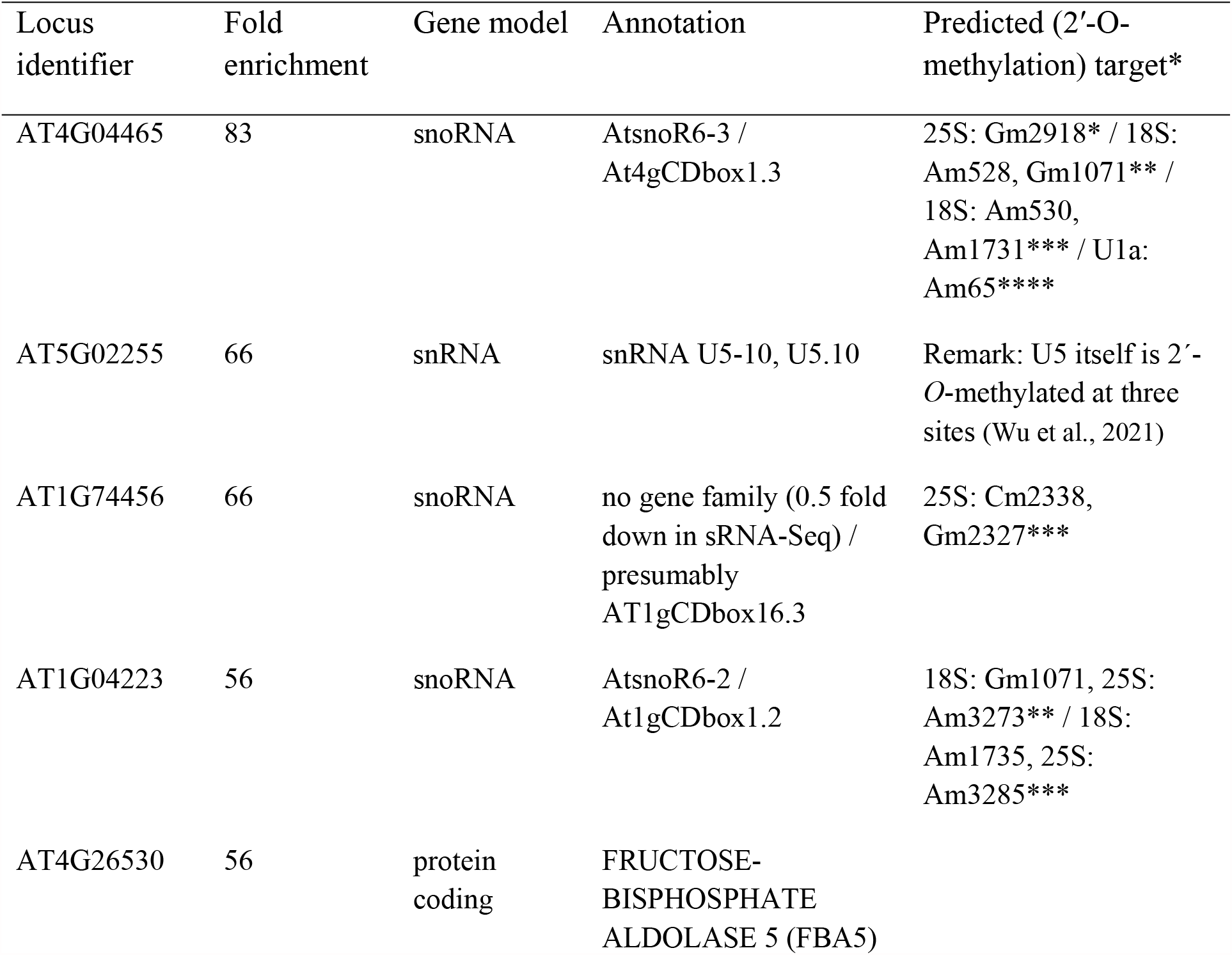
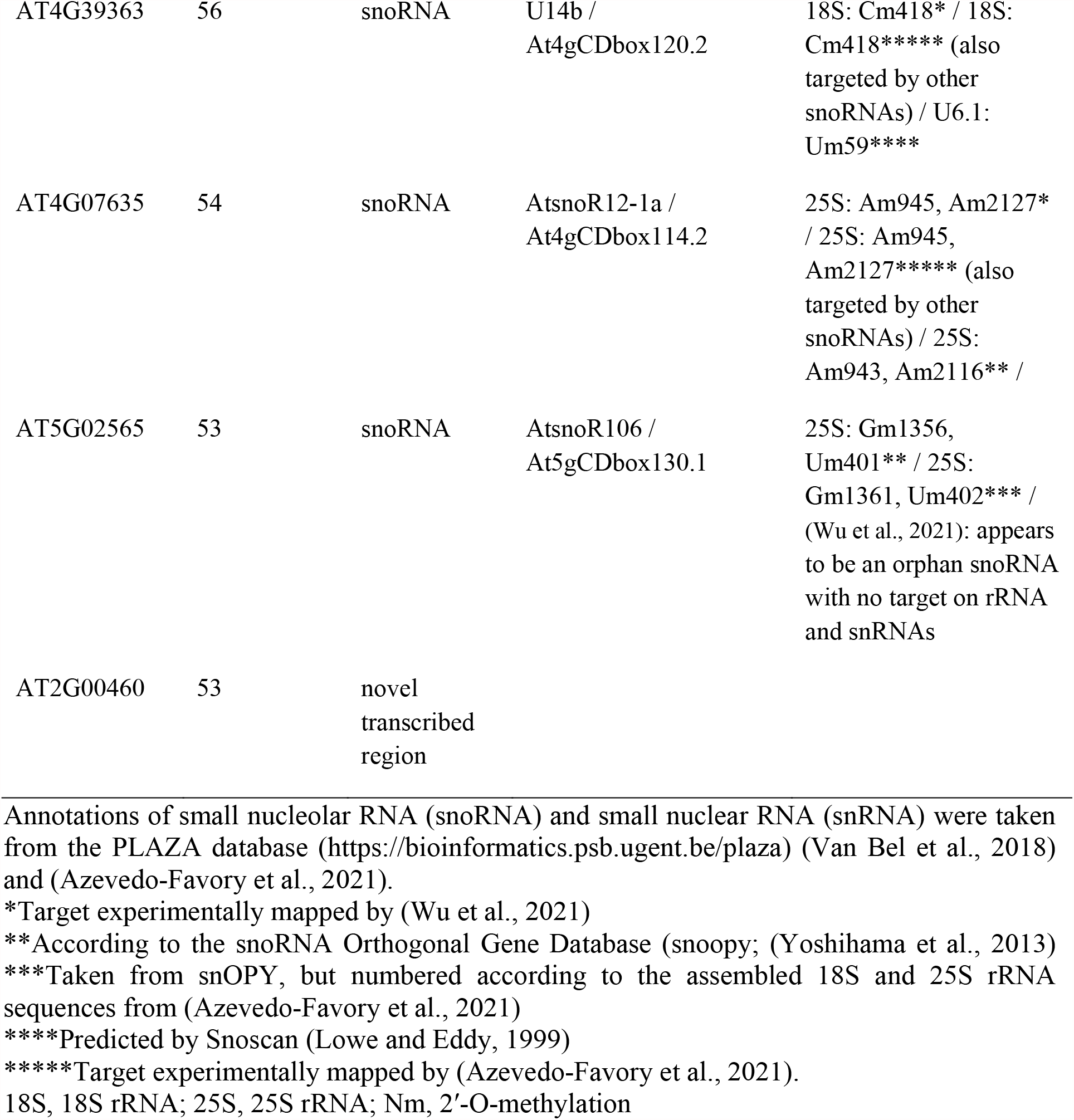
Top-ranking loci (> 50-fold enrichment) identified in RBP45D RIP-Seq experiments.

| Locus identifier | Fold enrichment | Gene model | Annotation | Predicted (2'-O-methylation) target* |
| --- | --- | --- | --- | --- |
| AT4G04465 | 83 | snoRNA | AtsnoR6-3 / At4gCDbox1.3 | 25S: Gm2918* / 18S: Am528, Gm1071** / 18S: Am530, Am1731*** / U1a: Am65**** |
| AT5G02255 | 66 | snRNA | snRNA U5-10, U5.10 | Remark: U5 itself is 2'-O-methylated at three sites (Wu et al., 2021) |
| AT1G74456 | 66 | snoRNA | no gene family (0.5 fold down in sRNA-Seq) / presumably AT1gCDbox16.3 | 25S: Cm2338, Gm2327*** |
| AT1G04223 | 56 | snoRNA | AtsnoR6-2 / At1gCDbox1.2 | 18S: Gm1071, 25S: Am3273** / 18S: Am1735, 25S: Am3285*** |
| AT4G26530 | 56 | protein coding | FRUCTOSE-BISPHOSPHATE ALDOLASE 5 (FBA5) |  |
| AT4G39363 | 56 | snoRNA | U14b /<br>At4gCDbox120.2 | 18S: Cm418* / 18S:<br>Cm418***** (also<br>targeted by other<br>snoRNAs) / U6.1:<br>Um59**** |
| AT4G07635 | 54 | snoRNA | AtsnoR12-1a /<br>At4gCDbox114.2 | 25S: Am945, Am2127*<br>/ 25S: Am945,<br>Am2127***** (also<br>targeted by other<br>snoRNAs) / 25S:<br>Am943, Am2116** / |
| AT5G02565 | 53 | snoRNA | AtsnoR106 /<br>At5gCDbox130.1 | 25S: Gm1356,<br>Um401** / 25S:<br>Gm1361, Um402*** /<br>(Wu et al., 2021): appears<br>to be an orphan snoRNA<br>with no target on rRNA<br>and snRNAs |
| AT2G00460 | 53 | novel<br>transcribed<br>region |  |  |

Annotations of small nucleolar RNA (snoRNA) and small nuclear RNA (snRNA) were taken from the PLAZA database (https://bioinformatics.psb.ugent.be/plaza) (Van Bel et al., 2018) and (Azevedo-Favory et al., 2021).
\*Target experimentally mapped by (Wu et al., 2021)
\*\*According to the snoRNA Orthogonal Gene Database (snoopy; (Yoshihama et al., 2013)
\*\*\*Taken from snOPY, but numbered according to the assembled 18S and 25S rRNA sequences from (Azevedo-Favory et al., 2021)
\*\*\*\*Predicted by Snoscan (Lowe and Eddy, 1999)
\*\*\*\*\*Target experimentally mapped by (Azevedo-Favory et al., 2021).
18S, 18S rRNA; 25S, 25S rRNA; Nm, 2'-O-methylation

SnoRNAs are short non-coding RNAs involved in ribosomal and spliceosome functions (Bratkovic et al., 2020). Based on their characteristic nucleotide motifs and association with canonical partner proteins, snoRNAs are classified into either C*/*D- (SNORD) or H*/*ACA-box (SNORA) subfamilies (Bratkovic et al., 2020), respectively catalyzing 2’
s-O-ribose methylation (Nm) and pseudouridylation of their substrates, mainly ribosomal RNAs (rRNA) (Bergeron et al., 2020). The snoRNAs identified here are all C/D-box members. Prediction and experimental identification of their targets (Table 1; Azevedo-Favory et al., 2021; Lowe and Eddy, 1999; Van Bel et al., 2018; Wu et al., 2021; Yoshihama et al., 2013) has suggested an involvement of RBP45D in 18S and 25S rRNA Nm. However, reverse transcription at low dNTP levels, followed by PCR (RTL-P; Dong et al., 2012), suggested that Nm of A1731 and A1735 of 18S rRNA and A945 of 25S rRNA (targets of snoR6-3, snoR6-2 and snoR12-1a, respectively) was not significantly altered in *rpb-ko* and *rls6* mutants compared to their WT (Figure S6C,D), nor was the 18S or 25S rRNA maturation pattern (Figure S6E).

Taken together, the RIP-Seq experiment, together with RNA-Seq data (see above) and Nm and Northern-blot analyses, point to an association of RBP45D with snRNPs and snoRNAs, but it does not seem to be involved in rRNA 2’
s-O-ribose methylation or maturation, and the results still do not explain the role of RBP45D in RdDM.

### RBP45D promotes siRNA production originating from the *LHCB1*.*2*:*LUC* transgene

In the canonical RdDM pathway, dsRNAs are cleaved by DCL3 into 24-nucleotide (nt) small interfering RNAs (siRNAs) which are loaded onto AGO effector proteins, which ultimately recruit the DNMT to catalyze *de novo* DNA methylation (see Graphical Abstract; Erdmann and Picard, 2020; Gebert and Rosenkranz, 2015; Zhang et al., 2018). Remarkably, a cross between *prors1* (*LUC*) and *dcl3-1* did not restore LUC expression (see Figure S1), indicating that, in the *prors1* (*LUC*) system, siRNAs are generated independently of DCL3. *DCL4* was identified in our screen, but in the corresponding mutant (*rls5*) and the *prors1* (*LUC*) *dcl4-2* double mutant, LUC expression is only rudimentarily restored, and not at all in leaf veins (see Figures 1 and S1). Therefore, to investigate whether the de-repression of *LUC* in *rls6* was actually accompanied by altered siRNA accumulation, deep sequencing of small RNAs (sRNAs; 17-30 nt in size) isolated from Col-0 (*LUC*), *prors1* (*LUC*) and *rls6* plants was carried out. The overall abundance and size distribution of sRNAs were comparable in all genotypes, and sRNAs of 24 nucleotides represented the predominant size class (Figure 6C) in accordance with previous reports (Elvira-Matelot et al., 2016). Consequently, RBP45D does not have a broad impact on overall sRNA accumulation. To test whether mutation of RBP45D might influence endogenous accumulation of sRNA at specific loci, the sequencing data were mapped to the TAIR10 genome. In *rls6*, relative to *prors1* (*LUC*), sRNAs derived from 44 (260) gene loci were at least 2-fold reduced (elevated) (Figure 6D and Table S7). Of the 260 loci sensitive to RBP45D, 15, 46, 51 and 148, correspond to miRNAs, long non-coding (lnc) RNAs, pseudo/transposable-element genes (non PCG) and protein-coding genes (PCG), respectively (Figure 6D), confirming the importance of RBP45D in regulating the production of sRNAs at discrete loci. The non-PCG and PCG loci regulated by RBP45D produce small RNAs that are predominantly 24 nt long in all genotypes (Figure 6E). Moreover, while lncRNA loci that produce 21-nt siRNAs predominate over 24-nt siRNAs in Col-0 (*LUC*) and *prors1* (*LUC*), this correlation was inverted in *rls6*, in which 24-nt siRNAs outnumbered 21-nt siRNAs. Notably, 71% of the sRNAs up-regulated in *rls6* relative to *prors1* (*LUC*) were also up-regulated in comparison to Col-0 (*LUC*), confirming that RBP45D also has a function that is independent of the *prors1* (*LUC*) background.

When sRNAs were mapped to the *LUC* cassette in Col-0 (*LUC*) no sRNAs originating from this region were identified, while in *prors1* (*LUC*) primarily 24-nt siRNAs derived from the *LHCB1*.*2* promoter and the Nos terminator (but not the *LUC* coding sequence) strongly accumulated (Figure 6F). In comparison with *prors1* (*LUC*), this accumulation was drastically reduced in the *rls6* mutant (Figure 6F, G), as confirmed by RNA gel blotting (Figure 6H). Because of RBP45D’s association with snoRNAs, we suspected that it might be involved in 2’
s- O-methylation of RNAs (see above). Nm deposition, mediated by HEN1, can also occur at 3′- terminal nucleotides of microRNAs (miRNAs) and siRNAs (Li et al., 2005; see also Graphical Abstract). In *Arabidopsis hen1* mutants, levels of miRNAs are reduced and size heterogeneity increased owing to the addition of uridines at the 3′ ends of small RNAs in the absence of 2′- O-methylation. No ladder of siRNAs was obvious in the *rls6* mutant when *LHCB1*.*2*-promoter-derived siRNAs were probed on Northern blots (Figure 6H), and the remaining reads that mapped to the *LHCB1*.*2* promoter region showed a clear 24-nt peak (Figure 6G). Therefore, it is unlikely that RBP45D is involved in 2’-O-methylation of siRNAs.

In summary, the accumulation of predominantly 24-nt siRNAs in the *prors1* (*LUC*) system is independent of DCL3, and is abrogated in the *rls6* mutant in which RBP45D is defective. To date, DCL3 is the only protein in Arabidopsis known to produce 24-nt siRNAs, which strongly implies that RBP45D itself is involved in the production of transgene-derived 24-nt siRNAs.

## Discussion

This study describes a forward genetics screen that identified the RNA-binding protein RBP45D as a new factor in the transcriptional silencing of transgenes (TGS). RBP45D is required for CHH methylation (Figure 2) and siRNA accumulation (Figure 6) of the *LHCB1*.*2* promoter/5′ UTR region of the transgenic *LHCB1*.*2*:*LUC* construct in the *prors1* (*LUC*) mutant. Genome-wide DNA methylation analysis of an *rbp*-*ko* line in the Col-0 background – independently of the *prors1* (*LUC*) system – showed that RBP45D also affects global DNA methylation, with particular emphasis on the stimulation of CHH methylation of transposon elements (TEs) and promoter regions (Figure 3). The RNA-directed DNA methylation (RdDM) pathway is involved in basic heat-stress tolerance (Popova et al., 2013), and a growing body of evidence has indicated that DNA methylation plays an important role in determining flowering times. For example, loss of transposon methylation in the *FLOWERING WAGENINGEN* (*FWA*) promoter activates the expression of *FWA*, which results in a late-flowering phenotype observed in *fwa* epigenetic alleles (Soppe et al., 2000). Lack of RBP45D also results in a late-flowering phenotype, especially at elevated temperatures (Figure 4). The *FWA* promoter was not hypomethylated in the *rbp-ko1* line (Table S3), but differential methylation of several genes affecting *FLC* mRNA expression, and the proposed role of FLC as a repressor of the thermal induction of flowering time (Gan et al., 2014), suggests that the late flowering phenotype of *rbp*-*ko* lines is provoked by elevated levels of FLC caused by epigenetic changes.

### Function of RNA-binding proteins associated with the spliceosome in RdDM

Another system that has been used to identify (anti)-silencing components employs the 35S- SUC2 line (Wang et al., 2013). When grown on a sucrose-containing medium, *35S-SUC2* transgenic plants over-accumulate sucrose because high expression of the SUCROSE-PROTON SYMPORTER 2 (SUC2) results in severe inhibition of root growth. Mutation of *ANTI-SILENCING 1* (*ASI1*) abrogates inhibition of root growth. ASI1 influences the RdDM rather indirectly. It is required for the production of full-length transcripts, as it stimulates distal polyadenylation downstream of intronic heterochromatic elements, including that found in the gene for INCREASE IN BONSAI METHYLATION 1 (IBM1) (Wang et al., 2013). IBM1 is a histone demethylase that is involved in the regulation of RdDM via the epigenetic control of *RDR2* and *DCL3* expression (Fan et al., 2012).

RBP45D is one of a family of eight proteins that show homology to yeast Nam8p, a non-essential U1 small nuclear ribonucleotide protein (snRNP) involved in the selection of weak 5′ splice sites (Puig et al., 1999). Our RIP-Seq analysis confirmed the association of

RBP45D with snRNPs, primarily U5 and U2 snRNPs (Table S6). Because two members of the *A. thaliana* RBP family have been shown previously not to stimulate splicing at weak splice sites (Lorkovic et al., 2000), (Kanno et al., 2020) have suggested that mutation of RBP45D could contribute to a hyper-GFP phenotype via a mechanism other than splicing regulation, but they did not rule out that RBP45D might suppress splicing at both weak and strong splice sites. We also detected only minor changes in splice patterns when RBP45D is lacking (Table S5). Moreover, transcripts coding for components of the RdDM are slightly elevated, but correctly spliced in *rbp-ko* lines, which precludes an indirect effect of the protein on RdDM like that seen for ASI1 (Wang et al., 2013), at least under normal growth conditions.

Other RBPs associated with TGS were identified principally between 2012 and 2015, and are also components of the spliceosome (Ausin et al., 2012; Dou et al., 2013; Du et al., 2015; Huang et al., 2013; Wang et al., 2013; Zhang et al., 2013). ARGININE/SERINE-RICH 45 (SR45) is an essential splicing factor (Ali et al., 2007) and was identified in a screen of T-DNA mutants that exhibited a late-flowering phenotype due to hypomethylation of *FWA* (Ausin et al., 2012). Interestingly, the *sr45-1* mutant also shows a late flowering phenotype owing to increased expression of *FLOWERING LOCUS C* (*FLC*) (Ali et al., 2007), which may also account for late flowering of *rbp*-*ko* lines (see above). Splicing of RdDM components in *sr45- 1* was not investigated, so it cannot be ruled out that the methylation phenotype of this mutant is a secondary effect. Alternatively, because 5S, AtSN1 and siR02 siRNAs are depleted in *sr45- 1*, it was proposed that SR45 might have a novel function in siRNA processing (Ausin et al., 2012). The remaining four spliceosome components implicated in silencing have been identified by He and coworkers by screening for mutants that restored *LUC* expression from an *RD29A* promoter-driven luciferase reporter (*RD29A*-*LUC*) in a *ros1* (Dou et al., 2013; Huang et al., 2013; Zhang et al., 2013) or *ros1 ago4* (Du et al., 2015) background. A mutation in *STABILIZED 1* (*STA1*), which codes for a PRP6-like splicing factor, has a drastic influence on Pol IV-dependent siRNA accumulation comparable with that of the *nrpe1* mutant. Moreover, Pol V-dependent RNA transcripts are also somewhat reduced in the *sta1* mutant. Based on these results, and because STA1 seems not to influence splicing of RdDM genes (as determined by RT-PCR), it was proposed that STA1 acts downstream of siRNA biogenesis and facilitates the production of Pol V-dependent RNA transcripts (Dou et al., 2013). Unlike the *sr45* and *sta1* mutants, mutations in RDM16 (pre-mRNA-splicing factor 3) (Huang et al., 2013) or PRP31 (Du et al., 2015) do not affect small RNA levels. In the *rdm16* mutant, DNA methylation is affected and levels of Pol V transcripts are decreased (Huang et al., 2013). Although RDM16 is involved in more than 300 intron-retention events, it does not affect the pre-mRNA splicing of known RdDM genes. Remarkably, only 15 genes show intron retention under normal growth conditions in *prp31*, but levels of mis-spliced mRNAs are markedly elevated under cold stress (Du et al., 2015). The scenario of stress-induced mis-splicing might also hold for *rbp-ko* lines under heat stress, but this remains to be tested. In the *zop1* (*zinc finger and OCRE domain-containing protein 1*) mutant, Pol IV-dependent siRNA accumulation and DNA methylation are reduced, but Pol V-dependent scaffold non-coding RNAs are not affected (Zhang et al., 2013). It is noteworthy that although PRP31, ZOP1, STA1 and RDM16 affect different steps in transcriptional gene silencing, at least PRP31, ZOP1 and STA1 interact physically (Du et al., 2015).

In the yeast *Cryptococcus neoformans*, stalled spliceosomes were identified as a signal for siRNA accumulation (Dumesic et al., 2013). However, the target transcripts are characterized by sub-optimal introns. The siRNAs identified in *prors1* (*LUC*) do not originate from intron-containing RNAs, thus excluding this possibility. In *A. thaliana*, spliceosome-associated proteins act at different levels on TGS, and details of how they participate in TGS remain unclear. But it is worth mentioning that the nuclear cap-binding complex, which is involved in pre-mRNA splicing, has a role in the DICER-LIKE1-dependent micro RNA pathway (Laubinger et al., 2008). This suggests that some components of the spliceosome, including RBP45D, may indeed play a direct role in the accumulation of specific siRNAs as well.

### A DCL3-independent mechanism for siRNA generation

In the canonical RdDM pathway, DCL3 cleaves dsRNAs generated by RDR2 into 24-nt siRNAs (Erdmann and Picard, 2020). A special feature of the *prors1* (*LUC*) system is that the accumulation of siRNAs originating from the transgenic *LHCB1*.*2* promoter is independent of functional DCL3 (Figure S1 and Graphical Abstract). DCL3 came to notice when it was found that some T-DNA insertion mutant lines induce 35S promoter homology-dependent TGS in its absence (Mlotshwa et al., 2010). The fact that a small pool of 24-nt siRNAs is still detectable in the *dcl2 3 4* triple mutant corroborates the assumption that at least some 24-nt siRNAs might be produced by as-yet unidentified factors (Yang et al., 2016; Ye et al., 2016; Zhang and Zhu, 2017).

Introduction of a *DCL4* mutation into *prors1* (*LUC*) reverses *LUC* silencing to some extent (Figure S1). DCL4 processes RDR6-generated dsRNA or Pol-II-derived hairpin-like structures into 21-nt siRNAs (Erdmann and Picard, 2020). Moreover, Ye et al. (2016) suggested that generation of DCL-independent siRNAs (sidRNAs) requires the action of AGO4 and RDR6. However, crossing of *prors1* (*LUC*) with *rdr6-15* did not mitigate *LUC* silencing either (Figure S1); therefore *LHCB1*.*2*-promoter-derived siRNAs might not fall into the class of sidRNAs.

How then are 24-nt siRNAs generated or stabilized independently of DCL3 in the *prors1* (*LUC*) system? Because RBP45D binds to C/D box snoRNAs that are involved in 2′-O- methylation of their RNA targets, we suspected that RBP45D might participate in a HEN1-like mechanism that stabilizes siRNAs by 2′-O-methylation of their 3′-termini. But the appearance of a clear 24-nt peak of reads mapping to the *LHCB1*.*2* promoter region argues against this hypothesis (see Figure 6G). Although it does not possess a double-stranded RNA-binding domain, it is tempting to speculate that RBP45D is at least part of the elusive DCL3- independent pathway (Zhang et al., 2018). Even if RBP45D might not be a direct slicer, it could help to stabilize either the precursor RNA for the subsequent cutting step, or the unidentified protein that performs the actual slicing step (Graphical Abstract). Alternatively, RBP45D could participate, as suggested for RRP6L1, in Pol IV-dependent siRNA production that may involve retention of Pol IV transcripts and/or in Pol V-produced lncRNA retention in chromatin to enable their scaffold function (Zhang et al., 2014). Finally, mutation of RBP45D prevents transgene silencing, which should facilitate the integration of properly expressed transgenes into plants.

## Acknowledgements

We thank Elisabeth Gerick for excellent technical assistance and Paul Hardy for critical reading of the manuscript. This work was supported by the Deutsche Forschungsgemeinschaft (TRR175, Project C01 to T.K. and Project C05 to D.L.).

## Author contributions

Conceptualization, T.K. and D.L.; Methodology, L.W., D.X., K.S., and T.K.; Validation, D.X.; Formal Analysis, T.K.; Investigation, L.W., D.X., K.S., and T.K.; Resources, D.L. and T.K.; Writing – Original Draft, T.K.; Writing – Review & Editing, L.W., D.X., D.L., W.F., and T.K.; Funding Acquisition, D.L. and T.K.; Supervision, T.K.

## Graphical abstract. Potential RBP45D action sites in DCL3-independen RNA-directed DNA methylation (RdDM)

A schematic diagram depicting the RdDM pathway in plants, together with the essential steps in post-transcriptional silencing (PTGS). Introduction of a *DCL4* mutation into *prors1* (*LUC*) only partially restores LUC activity. RBP45D may help to stabilize either the precursor RNA for the subsequent cleavage step or the – as yet unidentified – protein that performs the actual slicing step. Alternatively, RBP45D could participate in Pol IV-dependent siRNA production possibly involving retention of Pol IV transcripts and/or in Pol V-dependent lncRNA retention in chromatin to enable their scaffold function. Introduction of mutations shown in orange into *prors1* (*LUC*) reversed silencing; those in white did not, and those in grey were not tested.

## Methods

### Plant material and growth conditions

All seeds used in this study are in the Columbia-0 (Col-0) background, unless otherwise stated. T-DNA insertion mutants were obtained from the Nottingham Arabidopsis Stock Centre (NASC; http://arabidopsis.info/). Homozygous lines were identified using gene-specific primers and T-DNA border primers designed with the aid of the website http://signal.salk.edu/isectprimers.html. Details of T-DNA insertion sites in lines associated with RNA-directed DNA methylation can be found in (Wang et al., 2020b). Seeds were surface-sterilized with 0.6% (v/v) sodium hypochlorite solution containing 0.01% (v/v) Triton X-100 for 10 min, and then washed four times with ddH_2_O. After stratification at 4°C for 2 days, seeds were grown under long-day conditions (LD; 16 h light [100 μM of photons m^-2^ s^-1^]/8 h dark) at 22C for 25 days on soil or 1/2 Murashige and Skoog (MS) medium [with 0.5% (m/v) sucrose, 0.8% (m/v) plant agar], unless otherwise stated.

### 5-aza-2’-deoxycytidine (5Aza-dC) treatment and LUC activity assay

Prior to treatment of seedlings with 5Aza-dC (EMD Millipore, Cat. No. 189826), seeds were grown horizontally on 1/2 MS medium under LD for 3 days. The resulting seedlings were transferred to 1/2 MS medium supplemented with 50 μM 5Aza-dC and allowed to grow for another 7 days. These 2-week-old plants were then sprayed with 1 mM 5Aza-dC dissolved in 0.01% (v/v) Triton X-100 twice daily for another 7 days. For the detection of LUC activity, 1 mM luciferin (Thermo Fisher Scientific, Cat. No. L2916), dissolved in 0.01% (v/v) Triton X- 100, was sprayed on the surfaces of cotyledons or true leaves, which were then incubated in the dark for 5 min. Fluorescence images were taken immediately with a high-sensitivity CCD camera (Peqlab, Fusion Fx7).The exposure time was 5 min.

### Generation of *prors1* (*LUC*) plants

The region encompassing the 1-kbp promoter and the 5’UTR of *LHCB1*.*2* (*AT1T29910*) was amplified by PCR from genomic DNA with the primers described in Table S8, adding SalI and BamHI sites. After digestion with these enzymes, the product was inserted into the LUC^+^-NOS pPCV vector (Koncz et al., 1989). The construct was transformed into the *A. tumefaciens* strain GV3101, which was then introduced into Col-0 plants via floral dipping (Clough and Bent, 1998). Transgenic Col-0 (*LUC*) plants were selected on 1/2 MS medium supplemented with 50 μM hygromycin B (InvivoGen, Cat No. ant-hg-5). LUC activity was confirmed as described above. *prors1* (*LUC*) plants were generated by crossing homozygous Col-0 (*LUC*) with *prors1- 2* (GK-271H07).

### EMS mutagenesis of *Arabidopsis prors1* (*LUC*), screening and identification of second-site mutations

Mutagenesis of *Arabidopsis thaliana prors1* (*LUC*) seeds using ethyl methanesulfonate (EMS) was performed as described (Xu et al., 2020). Representative leaves from approximately 50,000 3-week-old M2 plants were painted with 1 mM luciferin (in 0.01% [v/v] Triton X-100) and LUC activity was checked as described above. Plants showing elevated LUC levels were classified as candidate suppressors of transgene silencing (*rls* mutants), and the phenotype was confirmed in the M3 generation.

To identify the causative mutations in *rls* mutants, they were backcrossed to the parental line *prors1 (LUC)*. F2 plants were first grown under LD conditions for 2 weeks before LUC activity was determined in representative leaves of each plant. Then 250 plants that showed increased LUC activity were selected from approximately 1,000 F2 plants. Leaves from each F2 population, as well as *prors1 (LUC)*, were pooled separately and ground in liquid nitrogen and genomic DNA was extracted as described (Wang et al., 2020a). About 2 μg of purified genomic DNA was used for next-generation sequencing, which was done as described before (Xu et al., 2020). For each sample, at least 7 GB of raw data were generated, which corresponds to more than 50-fold coverage of the *A. thaliana* genome, and single-nucleotide polymorphisms (SNPs) were identified as previously described (Xu et al., 2020). Only those SNPs that were supported by more than 4 reads with mapping quality >20 were kept. To identify the causative mutations in the *rls* mutants, the SNPs from each F2 population were compared with those from *prors1 (LUC)*, and the resulting *rls*-specific SNP lists were subjected to the web application CandiSNP (http://candisnp.tsl.ac.uk/), which generates SNP-density plots (Etherington et al., 2014). The output plots were screened first for the chromosome with the highest SNP density (with an allele frequency of >0.80), then for G/C to A/T transitions that were likely to be caused by EMS, and thirdly for non-synonymous amino-acid changes.

### Complementation of the *rls6* mutant and overexpression of RBP45D

For complementation of the *rls6* mutant, the gene *RBP45D* (*AT5G19350*) was amplified by PCR using primers listed in Table S8. The PCR product (without a stop codon) was inserted into the entry vector pDONR207 via the Gateway BP reaction, and subsequently cloned into the destination vector pGWB517 (35S pro, C-4xMyc) or pGWB541 (35S pro, C-EYFP) (Nakagawa et al., 2007) via the Gateway LR reaction (Invitrogen, Carlsbad, CA, USA). Both constructs were transformed into the *Agrobacterium tumefaciens* strain GV3101 and transferred to *rls6* mutant plants via floral dipping as described above. Transgenic plants were selected on 1/2 MS medium containing 50 μM hygromycin B as described above, and confirmed by LUC activity assay.

### Generation of CRISPR/Cas plants

The RBP45D-CRISPR/Cas vector was constructed as described previously (Wang et al., 2015). To select the optimal sgRNA, the program CHOP CHOP (http://chopchop.cbu.uib.no/) was used. Paired oligos (sgRNA-F and sgRNA-R; see Table S8) with 5′ overhangs were first treated with T4 polynucleotide kinase (NEB, M0201), then cooled from 95°C to 4°C at a 0.1°C/sec ramp rate. The annealed oligos were ligated to the pHEE401E vector via a GoldenGate reaction using the restriction enzyme BsaI (NEB, R3733) and T4 DNA ligase (NEB, M0202). The reaction was transformed directly into *E. coli* strain DH5α and positive colonies were identified on a LB plate containing kanamycin (50 µg/ml). The RBP45D-CRISPR/Cas vector was then transferred into the *Agrobacterium tumefaciens* strain GV3101, which was introduced into Col- 0 and *prors1(LUC)* plants via floral dipping as described above. Transgenic plants were first screened on 1/2 strength MS plates supplied with 50 μg/ml hygromycin. Putative DNA insertion/deletion sites in transgenic plants were then amplified using the paired oligos (CRISPR-iF and CRISPR-iR; see Table S8) and sequenced by the Sanger method.

### cDNA synthesis and quantitative RT-PCR analysis

Total RNA was extracted with the RNeasy Plant Mini kit (QIAGEN, Hilden, Germany) according to the manufacturer’s protocol, and 2 μg of the RNA was employed to synthesize cDNA using the iScript cDNA Synthesis Kit (Bio-Rad, Munich, Germany). Quantitative RT-qPCR analysis was performed on a Bio-Rad iQ5 real-time PCR instrument with the iQ SYBR Green Supermix (Bio-Rad, Munich, Germany) with the primers described in Table S8. Each sample was quantified in triplicate and normalized using *AT4G36800*, which codes for a RUB1- conjugating enzyme (RCE1), as an internal control.

### RNA gel-blot analysis

For gel-blot analysis of small RNAs, the protocol of (Pall and Hamilton, 2008) was modified. Total RNA was isolated with the TRIzol reagent (Invitrogen, Carlsbad, USA), and 10-µg aliquots were fractionated on a 15% polyacrylamide gel for 3 h at 60 V. The RNA gel was electroblotted onto an Amersham Hybond-N neutral membrane (GE Healthcare UK Limited, Buckinghamshire, UK) at 180 mA for 1 h. The RNA was fixed by using 1-ethyl-3-[3- dimethylaminopropyl] carbodiimide hydrochloride (EDC, Sigma-Aldrich) for 1 h at 60°C. Specific probes were designed that were complementary to the gene/sRNA of interest. Pre-hybridisation was done in Church buffer (1 M Na_2_HPO_4_, 1 M NaH_2_PO_4_, 0.5 M EDTA, pH 8.0, 20% SDS, 20 x SSC, 100x Denhardt’
ss reagent) for at least 2 h at the hybridisation temperature. For small Northern blots, DNA oligonucleotides (Table S8) were radiolabelled with [^32^P]γ-ATP (Hartmann Analytic GmbH, Braunschweig, Germany) using T4 kinase (New England Biolabs, Ipswich, MA, USA) according to the manufacturer’s protocol. Overnight hybridisation and washing was done according to (Pall and Hamilton, 2008). The signal was transferred to a phosphor imaging screen, which was scanned with the Typhoon TRIO (GE Healthcare Typhoon). The results were normalised to U6 by using the Quantity One Software (Bio-Rad, Hercules, CA, USA). Northern-blot detection of ribosomal RNAs was done as described previously (Romani et al., 2015).

### Reverse transcription at low dNTP followed by PCR (RTL-P)

irst-strand cDNA was synthesized from 500 ng of total RNA with 0.125 μg random RT primers. The mixture was heated to 70°C for 5 min, and the tube was immediately cooled on ice. The following components were added: (1) M-MLV reverse transcriptase (Promega, Cat. No. M1701), 1 mM dNTP mix, RNaseOUT (Invitrogen, Cat No. 10777019), and (2) the same conditions except for a 10-fold lower dNTP concentration (0.1 mM dNTP mix). These two reactions were incubated for 60 min at 37°C. qPCR was then performed in 10-μL reactions with SsoAdvanced Universal SYBR Green Supermix (Bio-Rad, Cat. No.1725271).

### Protein extraction and Western blot analysis

Protein extraction and Western blotting were performed as described before (Wang et al., 2020a). Approximately 100 mg of true leaves from 26-day-old transgenic plants were ground to a fine powder in liquid nitrogen. Proteins were extracted by adding 1 mL protein extraction buffer (20 mM HEPES [pH7.4]), 2 mM EDTA, 2 mM EGTA, 25 mM NaF, 1 mM Na_3_VO_4_, 50 mM glycerophosphate, 100 mM NaCl, 0.5% [v/v] Triton X-100, 10% [v/v] glycerol, 1x SIGMAFAST protease inhibitor), then mixing thoroughly mixed followed by incubation on ice for 30 min. Cell debris was removed by centrifugation at 12,000 *× g* for 30 min. Proteins in the supernatant (soluble fraction) were precipitated using 9 volumes of 100% acetone, 1/20 volume of 0.5 M Na_2_CO_3_ and 1/20 volume of 0.5 M DTT. After incubation at -20 °C for 30 min, proteins were pelleted by centrifugation at 3,000 *× g* for 30 min. The pellet was air-dried for 5 min at room temperature (RT) and dissolved in 200 μL of 1× Laemmli buffer. Proteins were fractionated on a 10% SDS-PA gel and transferred to PVDF membranes (Millipore, IPVH00010), which were then treated with 5% non-fat milk in 1× TBS buffer (50 mM Tris-HCl [pH 7.5] and 150 mM NaCl) for 1 h at RT. After overnight incubation at 4°C with antiserum against GFP (Sigma-Aldrich, G1546) or c-Myc (Santa Cruz Biotechnology, Dallas TX, USA, sc-40), the membrane was washed for 4 × 10 min with 1× TBST (50 mM Tris·HCl [pH 7.5], 150 mM NaCl, 0.1% Tween-20) at RT, the blot was incubated with goat-anti-rabbit or goat-anti-mouse IgG-HRP (Santa Cruz Biotechnology, sc-2004 or sc-2005) at RT for 1 h,and then washed for another 4 × 10 min with 1× TBST. Blots were sprayed with Pierce ECL Western Blotting Substrate (Thermo Fisher Scientific, Cat. No. 32106) and labelled bands were detected with a high-sensitivity CCD camera (Peqlab, Fusion Fx7).

### Confocal imaging

For confocal imaging, roots were excised from 5-day-old Arabidopsis seedlings grown vertically on 1/2 MS medium and immersed in 10 μM propidium iodide for 2 sec, followed by two 2-sec washes with ddH_2_O. Confocal pictures were taken with a Leica TCS SP5 laser-scanning confocal microscope. EYFP was excited with the 514-nm line of an argon laser, and the emission was collected by a 535/30 nm band-pass filter and was false colored green.

### Whole-genome bisulfite sequencing and analysis

A total amount of 5.2 µg genomic DNA spiked with 26 ng lambda DNA was fragmented by sonication to lengths of 200-400 bp with a Covaris S220 (Covaris, Woburn, MA, USA), followed by end-repair and adenylation. Cytosine-methylated barcodes were ligated to the sonicated DNA, and then these DNA fragments were treated twice with bisulfite using the EZ DNA Methylation-GoldTM kit (Zymo Research). The resulting single-stranded DNA fragments were PCR-amplified using KAPA HiFi HotStart Uracil + ReadyMix (2x). Library concentration was quantified with the Qubit® 2.0 Fluorometer (Life Technologies, Carlsbad, CA, USA) and by quantitative PCR, and the insert size was checked with the Agilent 2100 Bioanalyzer system (Agilent Technologies, Waldbronn, Germany). Library preparations were sequenced on an Illumina platform and 150-bp paired-end reads were generated (Novogene, Beijing, China). The Bismark software (Krueger and Andrews, 2011) was used to perform alignments against the TAIR10 genome with standard settings, and differential methylated regions (DMRs) were determined with an in-house (Novogene) script, or with MethylDackel (https://github.com/dpryan79/MethylDackel) and subsequent usage of Metilene (Juhling et al., 2016). Sequencing data have been deposited in NCBI’s Gene Expression Omnibus (Edgar et al., 2002) and are accessible through the GEO Series accession number xxx.

### mRNA sequencing (mRNA-Seq) and data analysis

Total RNA was isolated using Trizol (Invitrogen) and purified using Direct-zol™ RNA MiniPrep Plus columns (Zymo Research) according to the manufacturer’s instructions. RNA integrity and quality was assessed with an Agilent 2100 Bioanalyzer (Agilent, Santa Clara, USA). After the quality check, mRNA was enriched using oligo(dT) beads. Generation of RNA-Seq libraries and 150-bp paired-end sequencing on an Illumina HiSeq 2500 system (Illumina, San Diego, USA) were conducted at Novogene Biotech with standard Illumina protocols. Three independent biological replicates were used per genotype. RNA-Seq reads were analyzed on the Galaxy platform (Afgan et al., 2016) essentially as described before (Wang et al., 2020a), except that reads were mapped to the *Arabidopsis thaliana* genome (TAIR10) with the gapped-read mapper RNA STAR (Dobin et al., 2013) to detect splicing events. General splice site statistics were generated with QualiMap RNA-Seq QC (Okonechnikov et al., 2016), and reads counted with featureCounts (Liao et al., 2014) with the help of the gene annotation in Araport11 (www.araport.org/data/araport11) on the features “exon”, “protein”, and “CDS”, respectively. Intron retention events were detected by subtraction of the number of reads covering “CDS” from those covering “protein”. Differentially expressed genes were obtained with DESeq2 (Love et al., 2014), applying a 2- fold change cut-off and an adjusted *p* < 0.05. Sequencing data have been deposited in NCBI’s Gene Expression Omnibus (Edgar et al., 2002) and are accessible through the GEO Series accession number GSE176315.

### RNA immunoprecipitation and sequencing (RIP-Seq)

RNA immunoprecipitation (RIP) was performed as described previously (Terzi and Simpson, 2009) with small modifications, and all the buffers used were pre-chilled to 4°C. Basically, 14- day-old seedlings (*35S:RBP45D-c-Myc/rls6*) grown on 1/2 MS medium were harvested and washed for 4×1 min with cold (4°C) ddH_2_O. Then the seedlings were fixed with 1% formaldehyde (FA) for 20 min by vacuum infiltration at a pressure of 0.09 Mpa, and fixation was stopped by 5 mins of vacuum infiltration with 125 mM glycine. Fixed seedlings were then washed for 4×1 min with cold ddH_2_O, ground to a fine powder in liquid nitrogen and stored at -80°C. Around 400 mg of the powder was transferred to a pre-chilled 2-mL tube, 1.2 mL of RIP buffer (20 mM HEPES [pH 7.4], 2 mM EDTA, 2 mM EGTA, 25 mM NaF, 1 mM Na_3_VO_4_, 50 mM glycerophosphate, 500 mM NaCl, 10% glycerol, 0.5% Triton X-100, 1 mM PMSF, 1× protease inhibitor, 100 U/mL RNse inhibitor) was added, mixed and incubated on ice for 10 min. Samples were then centrifuged at 12,000 *×g* for 30 min at 4°C, and the resulting clear cell lysate was filtered through 0.45-μm filters. To prepare beads-antibody conjugate, 33 μL of Dynabeads Protein G slurry (Invitrogen, Cat. No. 10003D) was washed once with 200 μL of PBST (137 mM NaCl, 2.7 mM KCl, 10 mM Na_2_HPO_4_, 2 mM KH_2_PO_4_, 0.1% Tween-20) and incubated with 40 μL of the anti-Myc antibody (Santa Cruz Biotechnology, sc-40) in 200 μL PBST for 1 h at room temperature. For immunoprecipitation, 600 μl of the filtered cell lysate (the leftovers was reserved for immunoblot analysis and total RNA extraction) was added to the bead-antibody conjugate and incubated at 4°C for 4 h with slow agitation in the dark. To release the RNA, the beads were first washed four times for 5 min each with washing buffer (50 mM HEPES, 500 mM NaCl, 10% glycerol, 0.05% Triton X-100), and were then resuspended in 100 μL protease solution (30 mM Tris-HCl, pH 8.0, 50 µg/ml proteinase K) and incubated at 37°C for 30 min with low-speed agitation. The beads were then removed with a magnet, and the supernatant (100 μL) was transferred to a new 1.5-mL tube. To extract the released RNA, 300 μL ddH_2_O and 400 μL Trizol reagent (Life Technologies) were added to the supernatant, mixed well and incubated at 55°C for 5 min with high-speed agitation (1400 rpm). Then, 100 μL of chloroform-isoamyl alcohol (24:1) was added to each sample, mixed and centrifuged at 12,000 × *g* at 4°C for 15 min. Then 1 μL of 15 mg/ml GlycoBlue (Invitrogen, Cat No. AM9516), 1 volume of isopropanol and 1/10 volume of 3 M NaOAC (pH5.5) were added to the upper phase, mixed well and incubated at -20°C for 2 h. After centrifugation at 12,000 × *g* at 4°C for 30 min, the pellet was washed with 80% ethanol (pre-chilled at -20 °C) and resuspended in 10 μL RNase-free water. DNA contamination was removed using DNaseI (NEB, M0303S), Ribosomal RNA depletion and library preparation were achieved using the RiboMinus Plant Kit for RNA-Seq (Invitrogen, A10838-08) and the NEBNext Ultra II RNA Library Prep kit (E7775S), respectively. The RIP libraries were sequenced at Novogene Biotech on an Illumina HiSeq2500 system, and the RNA-Seq reads were analyzed on the Galaxy platform as described above.

### Preparation, sequencing and analysis of a small-RNA library

Total RNA was extracted from 26-day-old plants grown under LD conditions using the Trizol reagent (Invitrogen, Carlsbad, USA) and dissolved in RNase-free H_2_O. For preparation of a small-RNA (sRNA) library, 30 μg RNA was mixed with an equal volume of 2*×* RPA loading buffer (10 ml of 98% formamide, 200 µl 2 mM EDTA, 10 mg bromophenol blue, 10 mg xylene cyanol) and denatured at 94°C for 3 min. RNA samples were fractionated on a 15% TBE-PA by electrophoresis at 120 V for 2 h at 4°C, and the gel was stained with 5 µl SYBR Gold (Invitrogen, S11494) for 5 min in 50 ml 1*×* TBE buffer (89 mM Tris base, 89 mM boric acid, 1 mM EDTA, pH 8.0). The gel band corresponding to the 17-30 nt sRNA fraction was cut out, placed in a low-bind 1.5-ml tube containing 500 μl of 0.3 M NaCl, and ground with a RNase-free pestle. Then the tube was incubated at -80°C for 30 min at 4°C overnight under slow rotation. The gel slurry was filtered through a Spin-X filter (Corning, Cat. No. 8162) by centrifugation at 20,000 *×g* for 1 min at 4°C. The flow-through was transferred to two 1.5-ml tubes (each containing 250 μl flow-through) and 625 µl ethanol, 25 µl NaAc (3 M, pH 5.0) and 1 µl glycogen (Invitrogen, Cat. 10814010) was added. After 4 h at -80°C, the sRNA was pelleted by centrifugation at 20,000 *×g* for 30 min at 4°C. The pellets were washed twice with 80% ethanol, and dissolved in 7 µl RNase-free H_2_O. Small RNA libraries were prepared with the NEBNext^®^ Small RNA library prep kit (E7300) according to the user’s guide. Libraries were sequenced in the 50-bp single-end mode on an Illumina HiSeq 2500 system at Novogene Biotech, using standard Illumina protocols. Three independent biological replicates were used per genotype. RNA-Seq reads were analyzed on the Galaxy platform (Afgan et al., 2016). Reads were processed essentially as above, with the exception that filtered reads (> 17 nucleotides) were mapped to the TAIR10 genome with Bowtie2 (Langmead and Salzberg, 2012), and mature microRNAs were identified with the miRBase (http://www.mirbase.org; Kozomara et al., 2019)). Sequencing data have been deposited in NCBI’s Gene Expression Omnibus (Edgar et al., 2002) and are accessible through the GEO Series accession number GSE176514.

### Further data analysis, statistical tests

One-way ANOVA was performed to determine statistical significances between genotypes (*P* < 0.05) followed by a Tukey’s test for differences of group means at a 95% confidence interval with SPSS Statistics 17.0.

## Supplemental tables not included in Supplemental information file

**Table S1. Single-nucleotide polymorphisms (SNPs) and transgene insertion sites in *prors1***

**(*LUC*)**. Related to Figure 1.

**(A)** SNPs were determined from *prors1* (*LUC*) and Col-0 (*LUC*), respectively, with the Col-0 TAIR10 genome as a reference. SNPs of Col-0 (*LUC*) were subtracted from those of *prors1* (*LUC*) to detect the SNPs specific for *prors1* (*LUC*).

**(B)** Genomic DNA from a pool of *prors1* (*LUC*) plants was subjected to 150-bp paired-end sequencing. Reads were then mapped to the TAIR10 genome with RNAStar, and chimeric fusions between the *A. thaliana* genome and the left border (LB) of the pPCV vector were detected.

**Table S3. Differential methylated regions (DMRs) in 26-day-old, LD-grown *rbp-ko-1* plants compared to Col-0 as determined by whole-genome bisulfite sequencing**. Related to Figure 3.

**Table S4. Genes and transposable elements whose transcript levels changed by more than 2-fold in both 26-day-old LD-grown *rbp-ko-1* and *-2* plants compared to Col-0**. Related to Figure 5.

**Table S5. Splicing and intron-retention analyses of mRNA-Seq data generated from 26- day-old, LD-grown *rbp-ko-1* and Col-0 plants**. Related to Figure 5.

**Table S6. RNA targets of RBP45D as determined by RIP (RNA immunoprecipitation)- Seq experiments**. Related to Table 1.

**Table S7. Small RNAs that are more than 2-fold differentially expressed in 26-day-old *rls6* mutant plants compared to *prors1* (*LUC*) or Col-0 (*LUC*), respectively**. Related to Figure 6.

## References

Afgan, E., Baker, D., van den Beek, M., Blankenberg, D., Bouvier, D., Cech, M., Chilton, J., Clements, D., Coraor, N., Eberhard, C., et al. (2016). The Galaxy platform for accessible, reproducible and collaborative biomedical analyses: 2016 update. Nucleic Acids Res 44, W3–W10.

Ali, G.S., Palusa, S.G., Golovkin, M., Prasad, J., Manley, J.L., and Reddy, A.S. (2007). Regulation of plant developmental processes by a novel splicing factor. PLoS One 2, e471.

Ausin, I., Greenberg, M.V., Li, C.F., and Jacobsen, S.E. (2012). The splicing factor SR45 affects the RNA-directed DNA methylation pathway in Arabidopsis. Epigenetics 7, 29–33.

Azevedo-Favory, J., Gaspin, C., Ayadi, L., Montacie, C., Marchand, V., Jobet, E., Rompais, M., Carapito, C., Motorin, Y., and Saez-Vasquez, J. (2021). Mapping rRNA 2’-O-methylations and identification of C/D snoRNAs in Arabidopsis thaliana plants. RNA Biol, 1-18.

Bach-Pages, M., Homma, F., Kourelis, J., Kaschani, F., Mohammed, S., Kaiser, M., van der Hoorn, R.A.L., Castello, A., and Preston, G.M. (2020). Discovering the RNA-Binding Proteome of Plant Leaves with an Improved RNA Interactome Capture Method. Biomolecules 10.

Bergeron, D., Fafard-Couture, E., and Scott, M.S. (2020). Small nucleolar RNAs: continuing identification of novel members and increasing diversity of their molecular mechanisms of action. Biochem Soc Trans 48, 645–656.

Blevins, T., Podicheti, R., Mishra, V., Marasco, M., Wang, J., Rusch, D., Tang, H., and Pikaard, C.S. (2015). Identification of Pol IV and RDR2-dependent precursors of 24 nt siRNAs guiding de novo DNA methylation in Arabidopsis. Elife 4, e09591.

Bratkovic, T., Bozic, J., and Rogelj, B. (2020). Functional diversity of small nucleolar RNAs. Nucleic Acids Res 48, 1627–1651.

Clough, S.J., and Bent, A.F. (1998). Floral dip: a simplified method for Agrobacterium-mediated transformation of Arabidopsis thaliana. Plant J. 16, 735–743.

Daxinger, L., Hunter, B., Sheikh, M., Jauvion, V., Gasciolli, V., Vaucheret, H., Matzke, M., and Furner, I. (2008). Unexpected silencing effects from T-DNA tags in Arabidopsis. Trends Plant Sci 13, 4–6.

Dobin, A., Davis, C.A., Schlesinger, F., Drenkow, J., Zaleski, C., Jha, S., Batut, P., Chaisson, M., and Gingeras, T.R. (2013). STAR: ultrafast universal RNA-seq aligner. Bioinformatics 29, 15–21.

Dong, Z.W., Shao, P., Diao, L.T., Zhou, H., Yu, C.H., and Qu, L.H. (2012). RTL-P: a sensitive approach for detecting sites of 2’-O-methylation in RNA molecules. Nucleic Acids Res 40, e157.

Dou, K., Huang, C.F., Ma, Z.Y., Zhang, C.J., Zhou, J.X., Huang, H.W., Cai, T., Tang, K., Zhu, J.K., and He, X.J. (2013). The PRP6-like splicing factor STA1 is involved in RNA-directed DNA methylation by facilitating the production of Pol V-dependent scaffold RNAs. Nucleic Acids Res 41, 8489–8502.

Du, J.L., Zhang, S.W., Huang, H.W., Cai, T., Li, L., Chen, S., and He, X.J. (2015). The Splicing Factor PRP31 Is Involved in Transcriptional Gene Silencing and Stress Response in Arabidopsis. Mol Plant 8, 1053–1068.

Dumesic, P.A., Natarajan, P., Chen, C., Drinnenberg, I.A., Schiller, B.J., Thompson, J., Moresco, J.J., Yates, J.R., 3rd, Bartel, D.P., and Madhani, H.D. (2013). Stalled spliceosomes are a signal for RNAi-mediated genome defense. Cell 152, 957–968.

Edgar, R., Domrachev, M., and Lash, A.E. (2002). Gene Expression Omnibus: NCBI gene expression and hybridization array data repository. Nucleic Acids Res 30, 207–210.

Elvira-Matelot, E., Hachet, M., Shamandi, N., Comella, P., Saez-Vasquez, J., Zytnicki, M., and Vaucheret, H. (2016). Arabidopsis RNASE THREE LIKE2 Modulates the Expression of Protein-Coding Genes via 24-Nucleotide Small Interfering RNA-Directed DNA Methylation. Plant Cell 28, 406–425.

Erdmann, R.M., and Picard, C.L. (2020). RNA-directed DNA Methylation. PLoS Genet 16, e1009034.

Espinas, N.A., Saze, H., and Saijo, Y. (2016). Epigenetic Control of Defense Signaling and Priming in Plants. Front Plant Sci 7, 1201.

Etherington, G.J., Monaghan, J., Zipfel, C., and MacLean, D. (2014). Mapping mutations in plant genomes with the user-friendly web application CandiSNP. Plant Methods 10, 41.

Fan, D., Dai, Y., Wang, X., Wang, Z., He, H., Yang, H., Cao, Y., Deng, X.W., and Ma, L. (2012). IBM1, a JmjC domain-containing histone demethylase, is involved in the regulation of RNA-directed DNA methylation through the epigenetic control of RDR2 and DCL3 expression in Arabidopsis. Nucleic Acids Res 40, 8905–8916.

Gan, E.S., Xu, Y., Wong, J.Y., Goh, J.G., Sun, B., Wee, W.Y., Huang, J., and Ito, T. (2014). Jumonji demethylases moderate precocious flowering at elevated temperature via regulation of FLC in Arabidopsis. Nat Commun 5, 5098.

Gebert, D., and Rosenkranz, D. (2015). RNA-based regulation of transposon expression. Wiley Interdiscip Rev RNA 6, 687–708.

Gong, Z., Morales-Ruiz, T., Ariza, R.R., Roldan-Arjona, T., David, L., and Zhu, J.K. (2002). ROS1, a repressor of transcriptional gene silencing in Arabidopsis, encodes a DNA glycosylase/lyase. Cell 111, 803–814.

He, X.J., Hsu, Y.F., Pontes, O., Zhu, J., Lu, J., Bressan, R.A., Pikaard, C., Wang, C.S., and Zhu, J.K. (2009). NRPD4, a protein related to the RPB4 subunit of RNA polymerase II, is a component of RNA polymerases IV and V and is required for RNA-directed DNA methylation. Genes Dev 23, 318–330.

Hooper, C.M., Castleden, I.R., Tanz, S.K., Aryamanesh, N., and Millar, A.H. (2017). SUBA4: the interactive data analysis centre for Arabidopsis subcellular protein locations. Nucleic Acids Res 45, D1064–D1074.

Huang, C.F., Miki, D., Tang, K., Zhou, H.R., Zheng, Z., Chen, W., Ma, Z.Y., Yang, L., Zhang, H., Liu, R., et al. (2013). A Pre-mRNA-splicing factor is required for RNA-directed DNA methylation in Arabidopsis. PLoS Genet 9, e1003779.

Huang, D.W., Sherman, B.T., Tan, Q., Collins, J.R., Alvord, W.G., Roayaei, J., Stephens, R., Baseler, M.W., Lane, H.C., and Lempicki, R.A. (2007). The DAVID Gene Functional Classification Tool: a novel biological module-centric algorithm to functionally analyze large gene lists. Genome Biol 8, R183.

Juhling, F., Kretzmer, H., Bernhart, S.H., Otto, C., Stadler, P.F., and Hoffmann, S. (2016). metilene: fast and sensitive calling of differentially methylated regions from bisulfite sequencing data. Genome Res 26, 256–262.

Kanno, T., Lin, W.D., Fu, J.L., Chang, C.L., Matzke, A.J.M., and Matzke, M. (2017). A Genetic Screen for Pre-mRNA Splicing Mutants of Arabidopsis thaliana Identifies Putative U1 snRNP Components RBM25 and PRP39a. Genetics 207, 1347–1359.

Kanno, T., Venhuizen, P., Wu, M.T., Chiou, P., Chang, C.L., Kalyna, M., Matzke, A.J.M., and Matzke, M. (2020). A Collection of Pre-mRNA Splicing Mutants in Arabidopsis thaliana. G3 (Bethesda) 10, 1983–1996.

Koncz, C., Martini, N., Mayerhofer, R., Koncz-Kalman, Z., Korber, H., Redei, G.P., and Schell, J. (1989). High-frequency T-DNA-mediated gene tagging in plants. Proc Natl Acad Sci U S A 86, 8467–8471.

Kozomara, A., Birgaoanu, M., and Griffiths-Jones, S. (2019). miRBase: from microRNA sequences to function. Nucleic Acids Res 47, D155–D162.

Krueger, F., and Andrews, S.R. (2011). Bismark: a flexible aligner and methylation caller for Bisulfite-Seq applications. Bioinformatics 27, 1571–1572.

Kuhn, J.M., Breton, G., and Schroeder, J.I. (2007). mRNA metabolism of flowering-time regulators in wild-type Arabidopsis revealed by a nuclear cap binding protein mutant, abh1. Plant J 50, 1049–1062.

Langmead, B., and Salzberg, S.L. (2012). Fast gapped-read alignment with Bowtie 2. Nat Methods 9, 357–359.

Laubinger, S., Sachsenberg, T., Zeller, G., Busch, W., Lohmann, J.U., Ratsch, G., and Weigel, D. (2008). Dual roles of the nuclear cap-binding complex and SERRATE in pre-mRNA splicing and microRNA processing in Arabidopsis thaliana. Proc Natl Acad Sci U S A 105, 8795–8800.

Lee, K., Leister, D., and Kleine, T. (2021). Arabidopsis Mitochondrial Transcription Termination Factor mTERF2 Promotes Splicing of Group IIB Introns. Cells 10.

Leister, D., and Kleine, T. (2016). Definition of a core module for the nuclear retrograde response to altered organellar gene expression identifies GLK overexpressors as gun mutants. Physiol Plant 157, 297–309.

Li, J., Yang, Z., Yu, B., Liu, J., and Chen, X. (2005). Methylation protects miRNAs and siRNAs from a 3’-end uridylation activity in Arabidopsis. Curr Biol 15, 1501–1507.

Liao, Y., Smyth, G.K., and Shi, W. (2014). FeatureCounts: an efficient general purpose program for assigning sequence reads to genomic features. Bioinformatics 30, 923–930.

Lorkovic, Z.J., Wieczorek Kirk, D.A., Klahre, U., Hemmings-Mieszczak, M., and Filipowicz, W. (2000). RBP45 and RBP47, two oligouridylate-specific hnRNP-like proteins interacting with poly(A)+ RNA in nuclei of plant cells. RNA 6, 1610–1624.

Love, M.I., Huber, W., and Anders, S. (2014). Moderated estimation of fold change and dispersion for RNA-seq data with DESeq2. Genome Biol. 15, 550.

Lowe, T.M., and Eddy, S.R. (1999). A computational screen for methylation guide snoRNAs in yeast. Science 283, 1168–1171.

Matzke, M.A., Kanno, T., and Matzke, A.J. (2015). RNA-Directed DNA Methylation: The Evolution of a Complex Epigenetic Pathway in Flowering Plants. Annu Rev Plant Biol 66, 243–267.

Matzke, M.A., Mette, M.F., and Matzke, A.J. (2000). Transgene silencing by the host genome defense: implications for the evolution of epigenetic control mechanisms in plants and vertebrates. Plant Mol Biol 43, 401–415.

Matzke, M.A., and Mosher, R.A. (2014). RNA-directed DNA methylation: an epigenetic pathway of increasing complexity. Nat Rev Genet 15, 394–408.

McWhite, C.D., Papoulas, O., Drew, K., Cox, R.M., June, V., Dong, O.X., Kwon, T., Wan, C., Salmi, M.L., Roux, S.J., et al. (2020). A Pan-plant Protein Complex Map Reveals Deep Conservation and Novel Assemblies. Cell 181, 460-474 e414.

Mlotshwa, S., Pruss, G.J., Gao, Z., Mgutshini, N.L., Li, J., Chen, X., Bowman, L.H., and Vance, V. (2010). Transcriptional silencing induced by Arabidopsis T-DNA mutants is associated with 35S promoter siRNAs and requires genes involved in siRNA-mediated chromatin silencing. Plant J 64, 699–704.

Nakagawa, T., Suzuki, T., Murata, S., Nakamura, S., Hino, T., Maeo, K., Tabata, R., Kawai, T., Tanaka, K., Niwa, Y., et al. (2007). Improved Gateway binary vectors: high-performance vectors for creation of fusion constructs in transgenic analysis of plants. Biosci Biotechnol Biochem 71, 2095–2100.

Okonechnikov, K., Conesa, A., and Garcia-Alcalde, F. (2016). Qualimap 2: advanced multi-sample quality control for high-throughput sequencing data. Bioinformatics 32, 292–294.

Pall, G.S., and Hamilton, A.J. (2008). Improved northern blot method for enhanced detection of small RNA. Nat Protoc 3, 1077–1084.

Peng, M., Cui, Y., Bi, Y.M., and Rothstein, S.J. (2006). AtMBD9: a protein with a methyl-CpG-binding domain regulates flowering time and shoot branching in Arabidopsis. Plant J 46, 282–296.

Pesaresi, P., Masiero, S., Eubel, H., Braun, H.P., Bhushan, S., Glaser, E., Salamini, F., and Leister, D. (2006). Nuclear photosynthetic gene expression is synergistically modulated by rates of protein synthesis in chloroplasts and mitochondria. Plant Cell 18, 970–991.

Pooggin, M.M. (2013). How can plant DNA viruses evade siRNA-directed DNA methylation and silencing? Int J Mol Sci 14, 15233–15259.

Popova, O.V., Dinh, H.Q., Aufsatz, W., and Jonak, C. (2013). The RdDM pathway is required for basal heat tolerance in Arabidopsis. Mol Plant 6, 396–410.

Puig, O., Gottschalk, A., Fabrizio, P., and Seraphin, B. (1999). Interaction of the U1 snRNP with nonconserved intronic sequences affects 5’ splice site selection. Genes Dev 13, 569–580.

Reichel, M., Liao, Y., Rettel, M., Ragan, C., Evers, M., Alleaume, A.M., Horos, R., Hentze, M.W., Preiss, T., and Millar, A.A. (2016). In Planta Determination of the mRNA-Binding Proteome of Arabidopsis Etiolated Seedlings. Plant Cell 28, 2435–2452.

Romani, I., Manavski, N., Morosetti, A., Tadini, L., Maier, S., Kuhn, K., Ruwe, H., Schmitz-Linneweber, C., Wanner, G., Leister, D., et al. (2015). A Member of the Arabidopsis Mitochondrial Transcription Termination Factor Family Is Required for Maturation of Chloroplast Transfer RNAIle(GAU). Plant Physiol 169, 627–646.

Sheldon, C.C., Burn, J.E., Perez, P.P., Metzger, J., Edwards, J.A., Peacock, W.J., and Dennis, E.S. (1999). The FLF MADS box gene: a repressor of flowering in Arabidopsis regulated by vernalization and methylation. Plant Cell 11, 445–458.

Singh, J., Mishra, V., Wang, F., Huang, H.Y., and Pikaard, C.S. (2019). Reaction Mechanisms of Pol IV, RDR2, and DCL3 Drive RNA Channeling in the siRNA-Directed DNA Methylation Pathway. Mol Cell 75, 576-589 e575.

Singh, J., and Pikaard, C.S. (2019). Reconstitution of siRNA Biogenesis In Vitro: Novel Reaction Mechanisms and RNA Channeling in the RNA-Directed DNA Methylation Pathway. Cold Spring Harb Symp Quant Biol 84, 195–201.

Soppe, W.J., Jacobsen, S.E., Alonso-Blanco, C., Jackson, J.P., Kakutani, T., Koornneef, M., and Peeters, A.J. (2000). The late flowering phenotype of fwa mutants is caused by gain-of-function epigenetic alleles of a homeodomain gene. Mol Cell 6, 791–802.

Stroud, H., Greenberg, M.V., Feng, S., Bernatavichute, Y.V., and Jacobsen, S.E. (2013). Comprehensive analysis of silencing mutants reveals complex regulation of the Arabidopsis methylome. Cell 152, 352–364.

Terzi, L.C., and Simpson, G.G. (2009). Arabidopsis RNA immunoprecipitation. Plant J. 59, 163–168.

Vaillant, I., Tutois, S., Cuvillier, C., Schubert, I., and Tourmente, S. (2007). Regulation of Arabidopsis thaliana 5S rRNA Genes. Plant Cell Physiol 48, 745–752.

Van Bel, M., Diels, T., Vancaester, E., Kreft, L., Botzki, A., Van de Peer, Y., Coppens, F., and Vandepoele, K. (2018). PLAZA 4.0: an integrative resource for functional, evolutionary and comparative plant genomics. Nucleic Acids Res 46, D1190–D1196.

Wang, B., Jin, S.H., Hu, H.Q., Sun, Y.G., Wang, Y.W., Han, P., and Hou, B.K. (2012). UGT87A2, an Arabidopsis glycosyltransferase, regulates flowering time via FLOWERING LOCUS C. New Phytol 194, 666–675.

Wang, L., Leister, D., Guan, L., Zheng, Y., Schneider, K., Lehmann, M., Apel, K., and Kleine, T. (2020a). The Arabidopsis SAFEGUARD1 suppresses singlet oxygen-induced stress responses by protecting grana margins. Proc Natl Acad Sci U S A 117, 6918–6927.

Wang, L., Leister, D., and Kleine, T. (2020b). Chloroplast development and genomes uncoupled signaling are independent of the RNA-directed DNA methylation pathway. Sci Rep 10, 15412.

Wang, X., Duan, C.G., Tang, K., Wang, B., Zhang, H., Lei, M., Lu, K., Mangrauthia, S.K., Wang, P., Zhu, G., et al. (2013). RNA-binding protein regulates plant DNA methylation by controlling mRNA processing at the intronic heterochromatin-containing gene IBM1. Proc Natl Acad Sci U S A 110, 15467–15472.

Wang, Z.P., Xing, H.L., Dong, L., Zhang, H.Y., Han, C.Y., Wang, X.C., and Chen, Q.J. (2015). Egg cell-specific promoter-controlled CRISPR/Cas9 efficiently generates homozygous mutants for multiple target genes in Arabidopsis in a single generation. Genome Biol 16, 144.

Wendte, J.M., and Pikaard, C.S. (2017). The RNAs of RNA-directed DNA methylation. Biochim Biophys Acta Gene Regul Mech 1860, 140–148.

Wu, S., Wang, Y., Wang, J., Li, X., Li, J., and Ye, K. (2021). Profiling of RNA ribose methylation in Arabidopsis thaliana. Nucleic Acids Res.

Xu, D., Dhiman, R., Garibay, A., Mock, H.P., Leister, D., and Kleine, T. (2020). Cellulose defects in the Arabidopsis secondary cell wall promote early chloroplast development. Plant J 101, 156–170.

Yang, D.L., Zhang, G., Tang, K., Li, J., Yang, L., Huang, H., Zhang, H., and Zhu, J.K. (2016). Dicer-independent RNA-directed DNA methylation in Arabidopsis. Cell Res 26, 1264.

Ye, R., Chen, Z., Lian, B., Rowley, M.J., Xia, N., Chai, J., Li, Y., He, X.J., Wierzbicki, A.T., and Qi, Y. (2016). A Dicer-Independent Route for Biogenesis of siRNAs that Direct DNA Methylation in Arabidopsis. Mol Cell 61, 222–235.

Yoshihama, M., Nakao, A., and Kenmochi, N. (2013). snOPY: a small nucleolar RNA orthological gene database. BMC Res Notes 6, 426.

Zhang, C.J., Zhou, J.X., Liu, J., Ma, Z.Y., Zhang, S.W., Dou, K., Huang, H.W., Cai, T., Liu, R., Zhu, J.K., et al. (2013). The splicing machinery promotes RNA-directed DNA methylation and transcriptional silencing in Arabidopsis. EMBO J 32, 1128–1140.

Zhang, H., Lang, Z., and Zhu, J.K. (2018). Dynamics and function of DNA methylation in plants. Nat Rev Mol Cell Biol 19, 489–506.

Zhang, H., Tang, K., Qian, W., Duan, C.G., Wang, B., Zhang, H., Wang, P., Zhu, X., Lang, Z., Yang, Y., et al. (2014). An Rrp6-like protein positively regulates noncoding RNA levels and DNA methylation in Arabidopsis. Mol Cell 54, 418–430.

Zhang, H., and van Nocker, S. (2002). The VERNALIZATION INDEPENDENCE 4 gene encodes a novel regulator of FLOWERING LOCUS C. Plant J 31, 663–673.

Zhang, H., and Zhu, J.K. (2017). New discoveries generate new questions about RNA-directed DNA methylation in Arabidopsis. Natl Sci Rev 4, 10–15.

Zhou, M., and Law, J.A. (2015). RNA Pol IV and V in gene silencing: Rebel polymerases evolving away from Pol II’s rules. Curr Opin Plant Biol 27, 154–164.

Zhu, J.K. (2009). Active DNA demethylation mediated by DNA glycosylases. Annu Rev Genet 43, 143–166.

