## Supplemental information for "The RNA-binding protein RBP45D of Arabidopis plays a role in epigenetic control of flowering time and DCL3-independent RNA-directed DNA methylation"

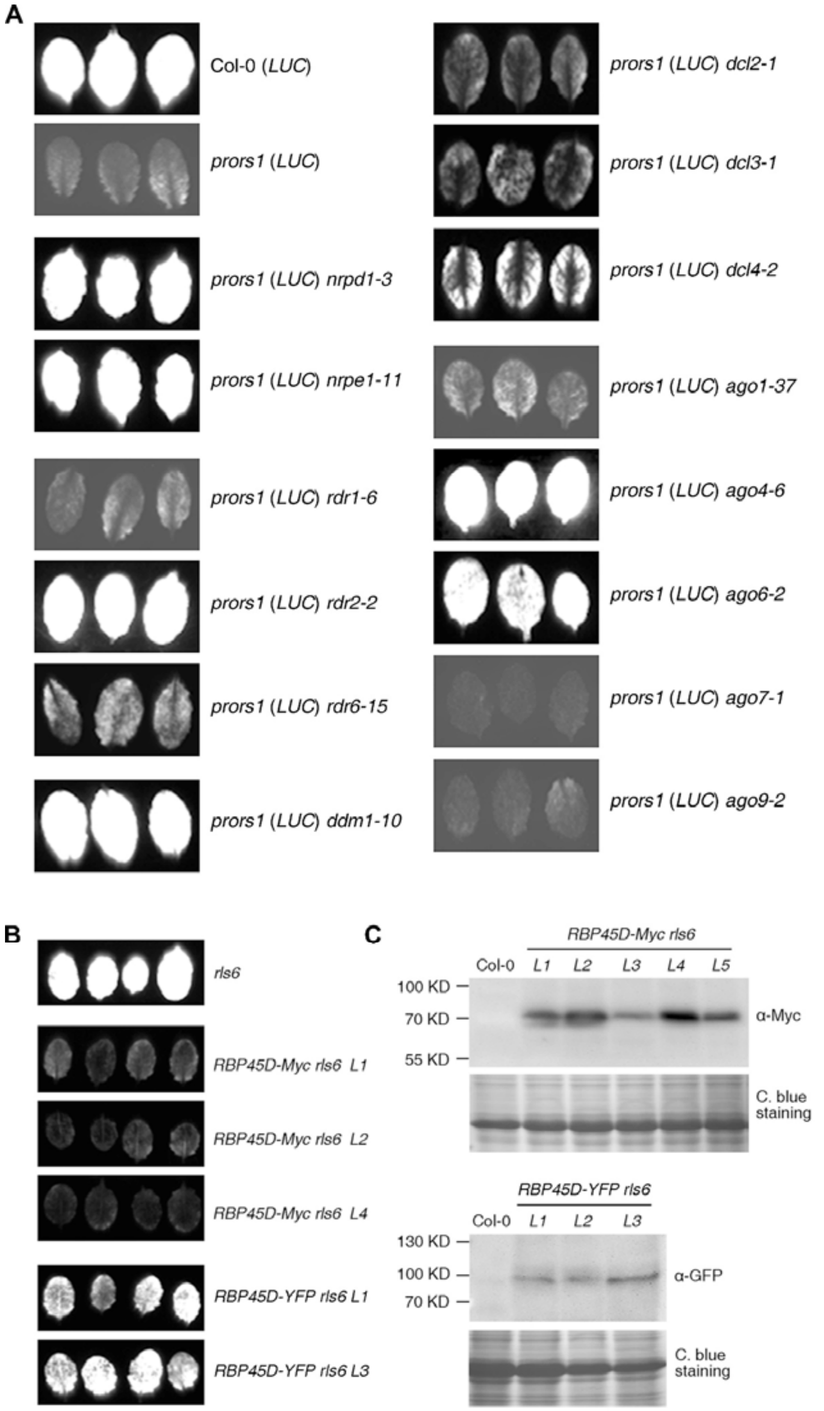


**Figure S1. Analysis of the function of diverse silencing components in the release of *LHCB1.2:LUC* silencing.** Related to Figure 1.

**(A)** Double mutants were generated by crossing *prors1 (LUC)* with various silencing mutants, and identified by genotyping with appropriate primers designed with the T-DNA Primer Design Tool (http://signal.salk.edu/tdnaprimers.2.html). Representative leaves from 3-week-old plants were sprayed with 1 mM luciferin and the chemical luminescence signals were captured using a highly sensitive CCD camera.

**(B)** Genetic complementation of *rls6* by *RBP45D*. Expression of an RBP45D-Myc fusion protein driven by the CaMV35S promoter fully restores the *prors1 (LUC)* phenotype, while RBP45D-YFP only does so partially.

**(C)** Detection of the expression of the RBP45D-Myc and -YFP fusion proteins by Western-blot analysis. C. blue, Coomassie Brilliant Blue; L, individual transgenic line.

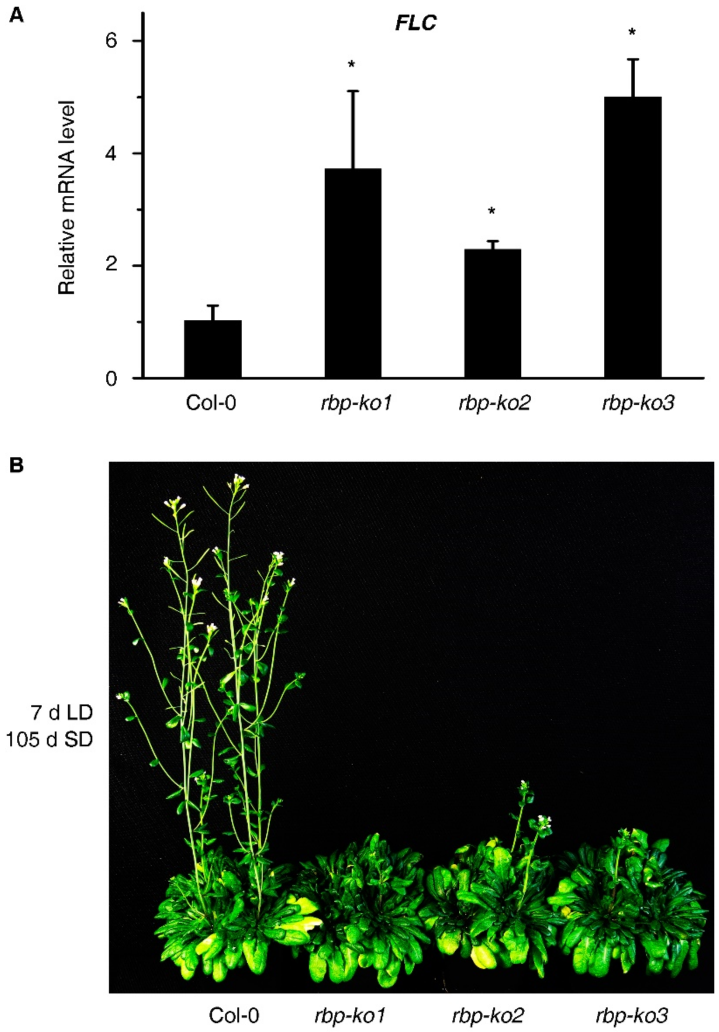


**Figure S2. Levels of *FLOWERING LOCUS C* (*FLC*) mRNA and the flowering phenotype of *rbp-ko* lines.** Related to Figure 4.

**(A)** Quantitative RT-PCR analysis of *FLC* mRNA levels in 26-day-old wild-type (Col-0) and *rbp-ko* plants. *AT4G36800*, which codes for an RUB1-conjugating enzyme (RCE1) served as the control. The RNA expression levels are reported relative to that in Col-0, which were set to 1. Data are represented as mean values from two independent experiments with three replicates each. Bars indicate standard deviations (SD). Significant differences between mutants and Col-0 control were evaluated with Student's t-test ($P < 0.05$) and are denoted by the asterisks (*).

**(B)** The *rbp-ko* lines exhibit a late flowering phenotype when grown under short-day (8 h light/16 h dark) conditions. Plants were first grown in LD conditions for 7 days and then transferred to SD conditions for 105 days.

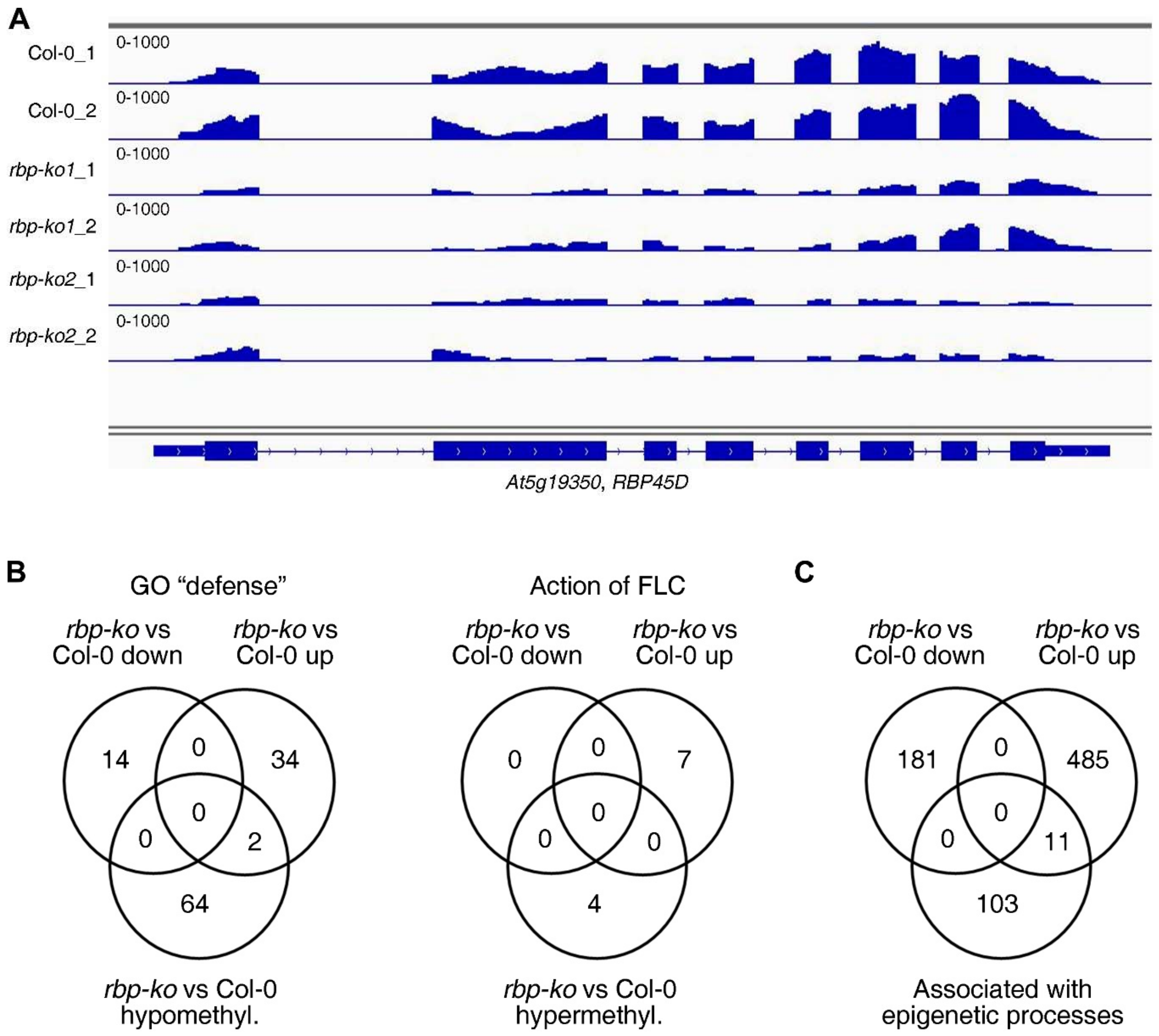


**Figure S3. Comparison of mRNA-Seq and WGBS data.** Related to Figures 3 and 5.
**(A)** Mapping of reads generated with mRNA-Seq of Col-0, *rbp-ko1* and *rbp-ko2* lines onto the *RBP45* gene. The read depths were visualized with the Integrative Genomics Viewer (IGV).
**(B)** Venn diagrams depicting the degree of overlap between the genes whose transcript levels changed at least 2-fold and those whose DNA methylation levels were altered in *rbp-ko* lines relative to Col-0.
**(C)** Venn diagram depicting the degree of overlap between the genes whose transcript levels changed at least 2-fold in *rbp-ko* lines compared to Col-0 and a set of genes coding for proteins involved in epigenetic processes.

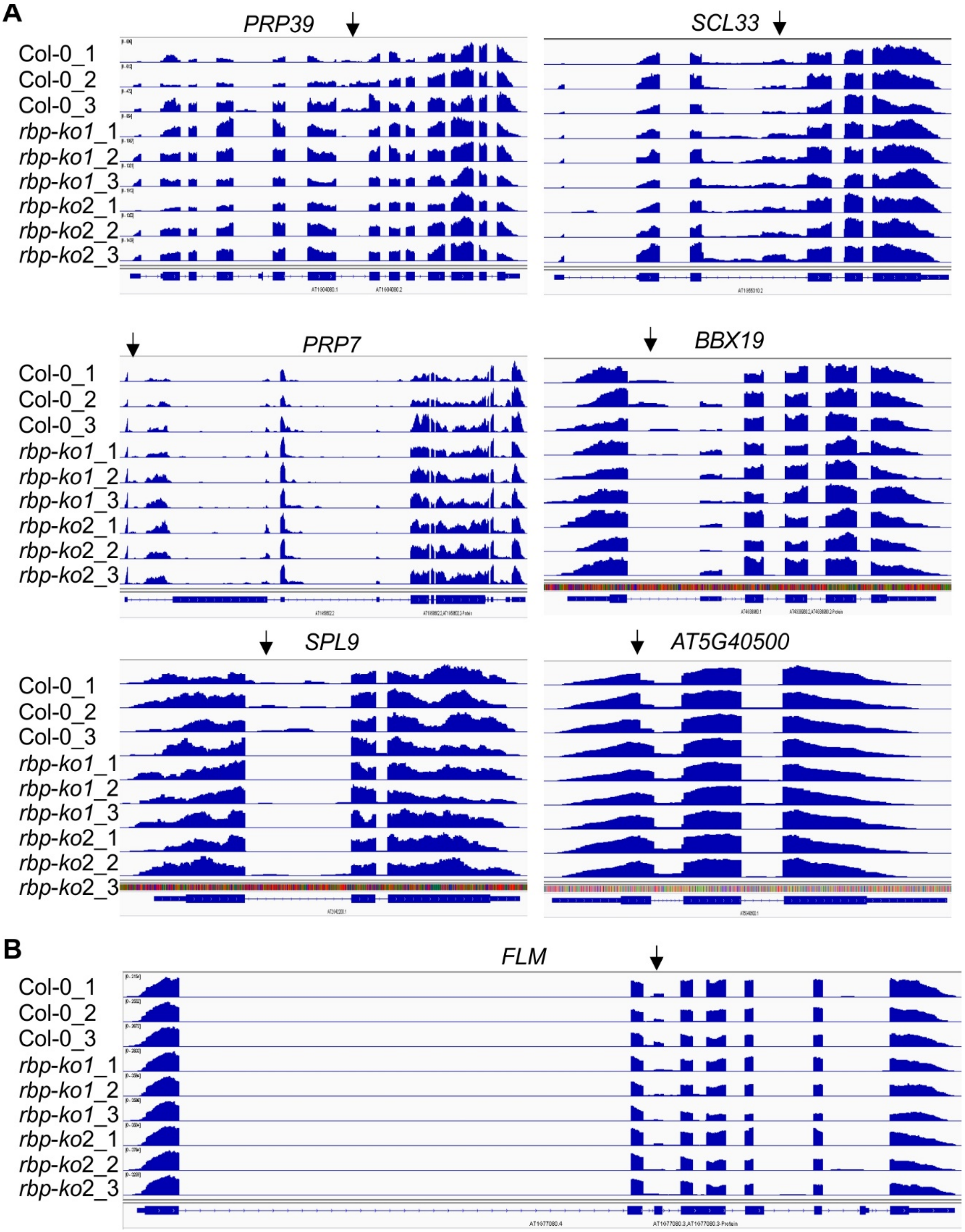


**Figure S4. Splicing patterns of transcripts coding for proteins related to splicing (PRP39 and SCL33), disease resistance (PRP7), flowering (FLM and BBX19), the vegetative to reproductive phase transition (SPL9) and a protein of unknown function (At5g40500).** Related to Figure 5.

RNA-Seq analysis was performed with RNA extracted from wild-type (Col-0) and *rbp-ko1* and -*2* mutant seedlings. The read depths were visualized with the IGV. The vertical arrows indicate differentially spliced regions.

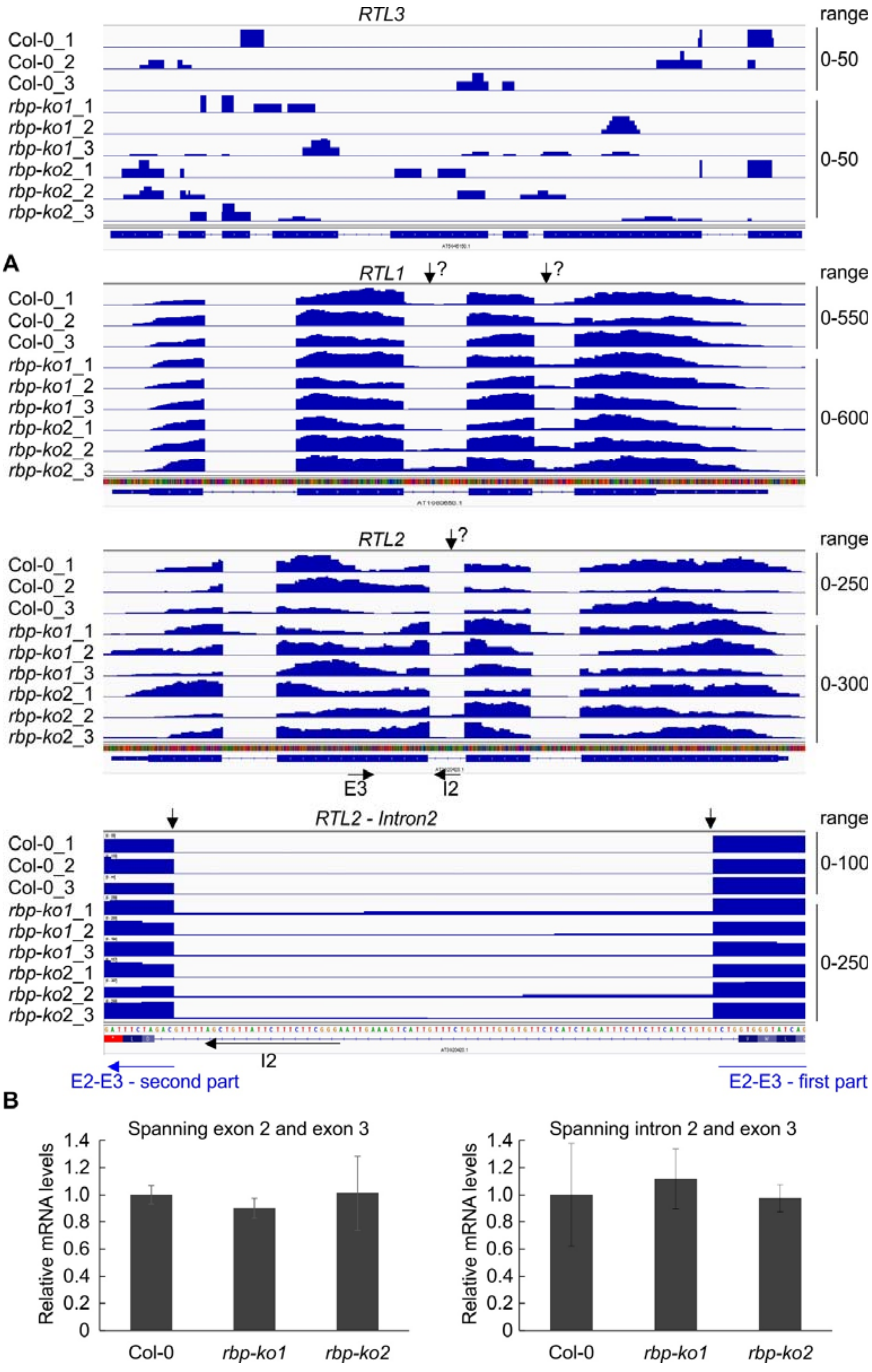


**Figure S5. Splicing patterns of transcripts coding for RNASE THREE-like proteins RTL1, RTL2 and RTL3.** Related to Figure 5.

**(A)** RNA-Seq analysis was performed with RNA extracted from wild-type (Col-0) and *rbp-ko1* and *-2* mutant seedlings. The read depths were visualized with the IGV. The vertical arrows indicate differentially spliced regions. Full-length transcripts of *RTL3* were below detection levels. The horizontal arrows indicate the primer positions used in panel **(B)**, the blue vertical disrupted arrow indicates the primer that spans exons 2 and 3. The vertical arrows with a question mark indicate putative differentially spliced regions.

**(B)** Quantitative RT-PCR analysis of 26-day-old wild-type (Col-0), *rbp-ko1* and *rbp-ko2*, and *prors* (*LUC*) and *rls6* mutant seedlings. PCR was conducted with the primer pairs marked in panel **(A)** and *AT4G36800* coding an RUB1-conjugating enzyme (RCE1) served as control. The RNA expression levels are reported relative to those in Col-0 (*rbp-ko1* and *rbp-ko2*), and *prors* (*LUC*) in the case of *rls6*. Data are presented as mean values from two independent experiments, each with three replicates. Bars indicate standard deviations (SDs). Significant differences between mutants and their respective control were evaluated with Student's t-test ($P < 0.05$) and are denoted by the asterisks (*).

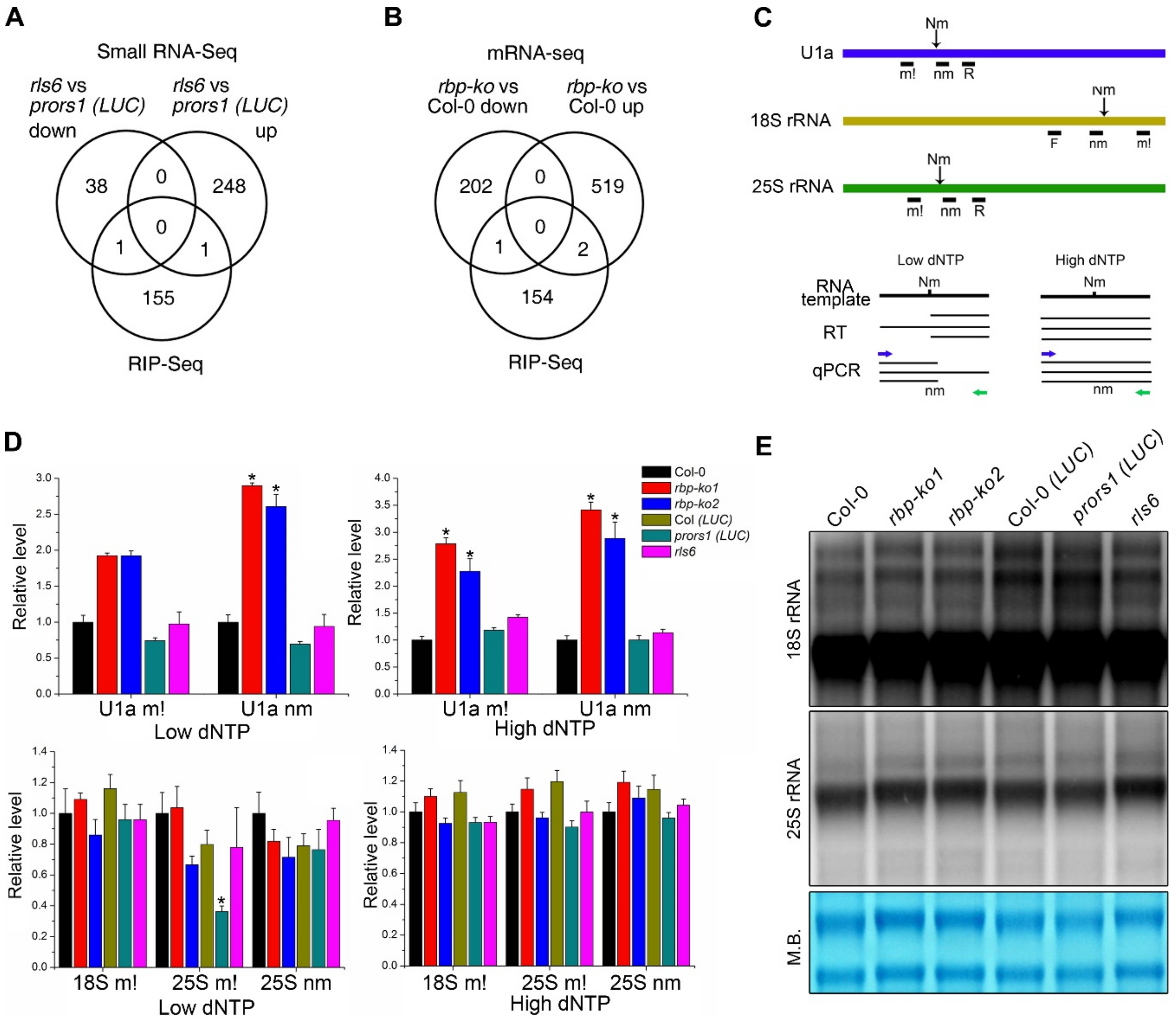


**Figure S6. RNAs bound by RBP45D do not overlap with the sRNAs or mRNAs regulated by RBP45D.** Related to Figures 5 and 6.

**(A)** Venn diagrams depicting the degree of overlap between the set of small RNAs altered by at least 2-fold in *rbp-ko* lines compared to Col-0 and the set of genes encoding RNAs that bind to RBP45D (RIP-Seq).

**(B)** Venn diagrams depicting the degree of overlap between genes whose transcript levels changed by at least 2-fold in *rbp-ko* lines compared to Col-0 and the set of genes that code for RNAs bound by RBP45D (RIP-Seq).

**(C, D)** Reverse transcription at low dNTP followed by PCR (RTL-P). Quantification of U1a and 18S and 25S ribosomal RNAs from each cDNA reaction (both high and low dNTP) was done by normalization to an internal reference gene (*RCE1*) to correct for differences in total RNA input and cDNA synthesis variations.

**(E)** Northern blot analysis depicting 18S and 25S rRNA maturation patterns.

**Table S2. Overview on mutations identified by the *prors1* (*LUC*) suppressor screen.** Related to Figure 1. Mutations were identified by whole-genome resequencing, confirmed by crossing *prors1* (*LUC*) plants with corresponding T-DNA lines and obtaining a high LUC phenotype and/or by complementation of the respective *rls* mutant.

| Mutant | Gene | Chr | Mutation | Crossed with T-DNA line | Complementation | Function |
|---|---|---|---|---|---|---|
| *rls1* | *RDR2* | 4 | $Arg_{258}$/Stop | Yes | Done/yes | RdDM pathway |
| *rls2* | *AGO4* | 2 | $Ser_{680}$/Phe | Yes | Done/yes | RdDM pathway |
| *rls3* | *AGO6* | 2 | $Gln_{210}$/Stop | Yes | Done/yes | RdDM pathway |
| *rls4* | *NRPD2A* | 3 | $Gly_{459}$/Arg | Yes | Not done | RdDM pathway |
| *rls5* | *DCL4* | 5 | $Gln_{362}$/Stop | Weak | Not done | RdDM pathway |
| *rls6* | *RBP45D* | 5 | $Ala_{193}$/Thr | Not done* | Done/yes | RNA binding protein, unknown function |

*Because the mutation is on the same chromosome like both the luciferase insertion and the T-DNA insertion into *PRORS1*, crossing was not used to confirm the identified locus. Pictures of restored LUC activity are represented in Fig. S1. Chr, chromosome; RdDM, RNA-directed DNA methylation; weak, only weak restored LUC activity; yes, high restored LUC activity.

**Table S8.** Primers used in this study.

| Atg number | Name | Primer sequence (from 5' to 3') |
|---|---|---|
| ***LHCB1.2-LUC* fusion** | | |
| *AT1G29910* | At1g29910SalI_s | TTACGTCGACTCAACAACACGAAAGAGT |
| | At1g29910BamHI_as | ATCGGGATCCTGAAACTTTTTGTGTTTT |
| **Primers for genotyping** | | |
| *AT5G19350* | GK-746D12 LP | TTTTGGAATTTCATCGAGTGG |
| | GK-746D12 RP | GGCATCTGTGTCCCATTGTAC |
| *AT5G19350* | SAIL_1304H10 LP | TTTACCTGTTGCGGATACTGG |
| | SAIL_1304H10 RP | CTGTTAGAGGTGCCAAGGTTG |
| **Complementation of the *rls6* mutant and overexpression of RBP45D** | | |
| *AT5G19350* | AT5G19350_attB1 | GGGGACAAGTTTGTACAAAAAAGCAGGCTAT GGCGATGATGCATCCTCCG |
| | AT5G19350_attB2 | GGGGACCACTTTGTACAAGAAAGCTGGGTGT TTGCCCAATTGTGATGTGAGCG |
| **Generation of CRISPR/Cas lines** | | |
| *AT5G19350* | sgRNA-F | ATTGCTTCTTCTGCAGAAGGCCT |
| | sgRNA-R | AAACAGGCCTTCTGCAGAAGAAG |
| | CRISPR-iF | GTCTGCTCATTTCTAGCTCCGT |
| | CRISPR-iR | AACCACATTCAGTGAACACAGC |
| **Real-time PCR** | | |
| *AT4G36800* | RCE1 RT F | CTGTTCACGGAACCCAATTC |
| | RCE1 RT R | GGAAAAAGGTCTGACCGACA |
| *AT5G19350* | RBP45D RT F | TCTTTGTTGGAGATTTAGCACCTG |
| | RBP45D RT R | ACTGGGTAAACAGCTTTGGT |
| *AT2G47580* | U1a m! F | TTGTGATCAATAAGACGAGTGG |
| | U1a nm F | GGGTGCTTAGCTTAAGGTCTC |
| | U1a R | CCCTCTGCCACAAATAATGAC |
| *18S rRNA* | 18S 1731 F | GTCGCTCCTACCGATTGAATG |
| | 18S 1731 m! R | CTCCTTCCTCTAAATGATAAGGT |
| | 18S 1731 nm R | GGACTTCTCGCGACGTCGCG |
| *25S rRNA* | 25S 945 m! F | GCGAAAGACTAATCGAACCATCTA |

| | | |
|---|---|---|
| | 25S 945 nm F | CTGGTTCCCTCCGAAGTTTCC |
| | 25S 945 R | AAGTTTGAGAATAGGTCGAGG |
| *AT3G20420* | RTL2_E2_E3_RT_F | AGACTATGGGTGATCTTTAGGG |
| | RTL2_E3_RT_R | GCCTTGCGATATCTTTATTCTCAG |
| | RTL2_I2_RT_F | AGGGCTTCTTTCTTATTGTCG |
| *LUC* | LUC_RT_F | CTGGAGAGCAACTGCATAAGGCTATG |
| | LUC_RT_R | CTATGTCTCCAGAATGTAGCC |
| *AT1G29910* | LHCB1.2 RT F | CCGTGAGCTAGAAGTTATCC |
| | LHCB1.2 RT R | GTTTCCCAAGTAATCGAGTCC |
| *AT5G10140* | FLC RT F | GGCAAGCTCTACAGCTTCTC |
| | FLC RT R | AACATGAGTTCGGTCTTCTTGG |
| **RNA gel blot analysis** | | |
| *AT1G29910* | LHCB1.2-sRNA | GCAGTCTCTAGTCCACTAGCTTTT |
| *18S rRNA* | 18S Nth F | TTAAAGGAATTGACGGAAGGG |
| | 18S Nth R | AACATCTAAGGGCATCACAGAC |
| *25S rRNA* | 25S 945 m! F | GCGAAAGACTAATCGAACCATCTA |
| | 25S 945 R | AAGTTTGAGAATAGGTCGAGG |
